# The latency of a visual evoked potential tracks the onset of decision making

**DOI:** 10.1101/275727

**Authors:** Michael D. Nunez, Aishwarya Gosai, Joachim Vandekerckhove, Ramesh Srinivasan

**Affiliations:** Department of Cognitive Sciences University of California Irvine CA USA; Department of Biomedical Engineering, University of California, Irvine, CA, USA; Massachusetts General Hospital and Harvard Medical School, Boston, MA, USA; Department of Statistics, University of California, Irvine, CA, USA; Institute of Mathematical Behavioral Sciences, University of California, Irvine, CA, USA

**Keywords:** decision making, figure-ground segregation, EEG, visual evoked potentials (VEP), diffusion models, hierarchical Bayesian methods

## Abstract

Encoding of a sensory stimulus is believed to be the first step in perceptual decision making. Previous research has shown that electrical signals recorded from the human brain track evidence accumulation during perceptual decision making (Gold and Shadlen, 2007; O’Connell et al., 2012; Philiastides et al., 2014). In this study we directly tested the hypothesis that the latency of the N200 recorded by EEG (a negative peak occurring between 150 and 275 ms after stimulus presentation in human participants) reflects the visual encoding time (VET) required for completion of figure-ground segregation before evidence accumulation. We show that N200 latencies vary across individuals, are modulated by external visual noise, and increase response time by *x* milliseconds when they increase by *x* milliseconds, reflecting a linear regression slope of 1. Simulations of cognitive decision-making theory show that variation in human response times not related to evidence accumulation (including VET) are tracked by the fastest response times. A relationship with a slope of 1 between N200 latencies and VET was found by fitting a linear model between trial-averaged N200 latencies and the 10th percentiles of response times. A slope of 1 was also found between single-trial N200 latencies and response times. Fitting a novel neuro-cognitive model of decision-making also yielded a slope of 1 between N200 latency and non-decision time in multiple visual noise conditions, indicating that N200 latencies track the completion of visual encoding and the onset of evidence accumulation. The N200 waveforms were localized to the cortical surface at distributed temporal and extrastriate locations, consistent with a distributed network engaged in figure-ground segregation of the target stimulus.

**Significance Statement:** Encoding of a sensory stimulus is believed to be the first step in perceptual decision making. In this study, we report evidence that visual evoked potentials (EPs) around 200 ms after stimulus presentation track the time of visual figure-ground segregation before the onset of evidence accumulation during decision making. These EP latencies vary across individuals, are modulated by external visual noise, and increase response time by *x* milliseconds when they increase by *x* milliseconds. Hierarchical Bayesian model-fitting was also used to relate these EPs to a specific cognitive parameter that tracks time related to visual encoding in a decision-making model of response time. This work adds to the growing literature that suggests that EEG signals can track the component cognitive processes of decision making.

## Introduction

We define *visual encoding time* (VET) as the amount of time for visual processing to occur in the human brain before decision processes can begin. While there has been abundant evidence of visual information appearing in the primary visual cortex approximately 60 milliseconds (ms) after stimulus onset (Schmolesky et al., 1998; Luck, 2012), further processing takes place within a network of brain areas comprising the visual system. The key objective of this further processing is the grouping together of elements of the scene coded in separate neurons into distinct objects, a process known as figure-ground segregation (Lamme, 1995). Masking, by presenting a distractor stimulus soon after the target stimulus onset, can interrupt figure-ground segregation up to 100 ms by degrading the encoding of the visual stimulus (Lamme et al., 2002). This work suggests that VET of the target stimulus is approximately 160 ms (100 ms masking time plus 60 ms for the time of the distractor stimulus to reach primary visual cortex). Furthermore, on the basis of ERP studies in humans and physiological studies in animal models (Bach and Meigen, 1992; Roitman and Shadlen, 2002; Straube et al., 2010), VET is expected to be 150 to 200 ms in duration. Our goals in the following study were to 1) clarify estimation of VET in human participants using EEG and 2) verify these time estimates in the framework of cognitive theory of quick human decision-making.

Human response times (time between stimulus onset and motor action execution) in decision making is theorized to be composed of VET, evidence accumulation time, response preparation time, and motor execution time. VET is separate from the decision-related processing, which has been experimentally found to reflect a sequential accumulation of evidence on level of the neuron (Gold and Shadlen, 2007; Shadlen and Kiani, 2013), the level of the scalp-recorded electroencephalograph (EEG; O’Connell et al., 2012; Philiastides et al., 2014; Kelly and O’Connell, 2015), and on the cognitive level as evidenced by cognitive models of human behavior (Voss et al., 2004; Ratcliff and McKoon, 2008). While evidence accumulation and motor planning may operate in parallel (Dmochowski and Norcia, 2015; Servant et al., 2015, 2016), it can be reasonably expected that figure-ground segregation is a sequential processing step that occurs before any evidence accumulation or motor planning. That is, the grouping of neural activity elicited by the stimulus across the visual system into a distinct “object” through figure-ground segregation is necessary before evidence accumulation and motor planning.

The figure-ground segregation speed of the human visual system has previously been estimated between 150 ms and 225 ms in humans with time-locked event related potentials (ERPs; Bach and Meigen, 1992; Thorpe et al., 1996; Vanrullen and Thorpe, 2001) using clever differences between experimental conditions. Loughnane et al. (2016) found initial evidence by looking at condition differences to suggest that stimulus-locked negative peaks around 200 ms (N200) influence the onset of a neural correlate of evidence accumulation. In other studies, single-trial EEG potentials thought to be generative of N200 peaks were suspected to be related to a “pre-attentive” phase before evidence accumulation (Zhang et al., 2016, 2018). Interestingly, single-neuron recordings of evidence accumulation from lateral intraparietal areas (LIP) in primates typically begin at similar time periods (*≈* 200 ms) after an experimental stimulus is displayed (Roitman and Shadlen, 2002; Shadlen and Kiani, 2013).

In this study, our hypothesis was that N200 peak-latencies would track VET no matter the external visual noise condition, participant, or participant’s internal state that changed across EEG sessions. Using both direct linear regressions between EEG latency and fast response times and novel neuro-cognitive models of decision making, we verified that N200 peak-latencies tracked non-decision time (NDT), of which visual encoding time (VET) is an additive component, with regression slopes that imply an *x* millisecond increase in N200 peak-latency leads to an *x* millisecond increase in NDT (i.e. a slope parameter of 1). We extend the theory of human perceptual decision-making by including this EEG-derived latency estimate of evidence accumulation onset.

## Materials and Methods

### Experiments and Participants

In order to evaluate the hypothesis that N200 latencies track VET, data was analyzed from two similar experiments. Data from Experiment 1 consisted of EEG recordings and behavioral observations from 12 unique participants (3 female, 9 male) with 2 sessions each. Data from Experiment 2 consisted of EEG recordings and behavioral observations from 4 unique participants (2 female, 2 male) with 7 sessions each. Two male participants participated in both experiments resulting in 14 unique participants across both experiments (5 female, 9 male) with 52 unique EEG sessions. Sessions of EEG collection and task performance for each participant in Experiment 1 were separated by at least 24 hours. Sessions of EEG collection and task performance for each participant in Experiment 2 were separated by 1 week. All participants had normal or corrected-to-normal vision and had no history or family history of epilepsy. All participants gave written informed consent, and all data was collected at the University of California, Irvine with approval by the Institutional Review Board.

### Experimental tasks

Data was included from two different experiments in this study in order to verify that any evidence for our hypothesis that we found (i.e. that N200 peak-latencies track VET) did not depend on the type of distracting visual stimuli. While we expect the distracting visual noise to yield different N200 peak-latencies and VETs across both experiments, we expect to find the same tracking of VET by the N200 peak-latency in both experiments across multiple visual contrast conditions. In both experiments, participants performed two-alternative forced choice tasks, specifically to determine whether a Gabor stimulus contained high or low spatial frequency content. Stimuli for Experiments 1 and 2 are built and displayed using the MATLAB Psychophysics toolbox (Psychtoolbox-3; www.psychtoolbox.org). Example stimuli are given in **Figure 1**.

**Figure 1.**
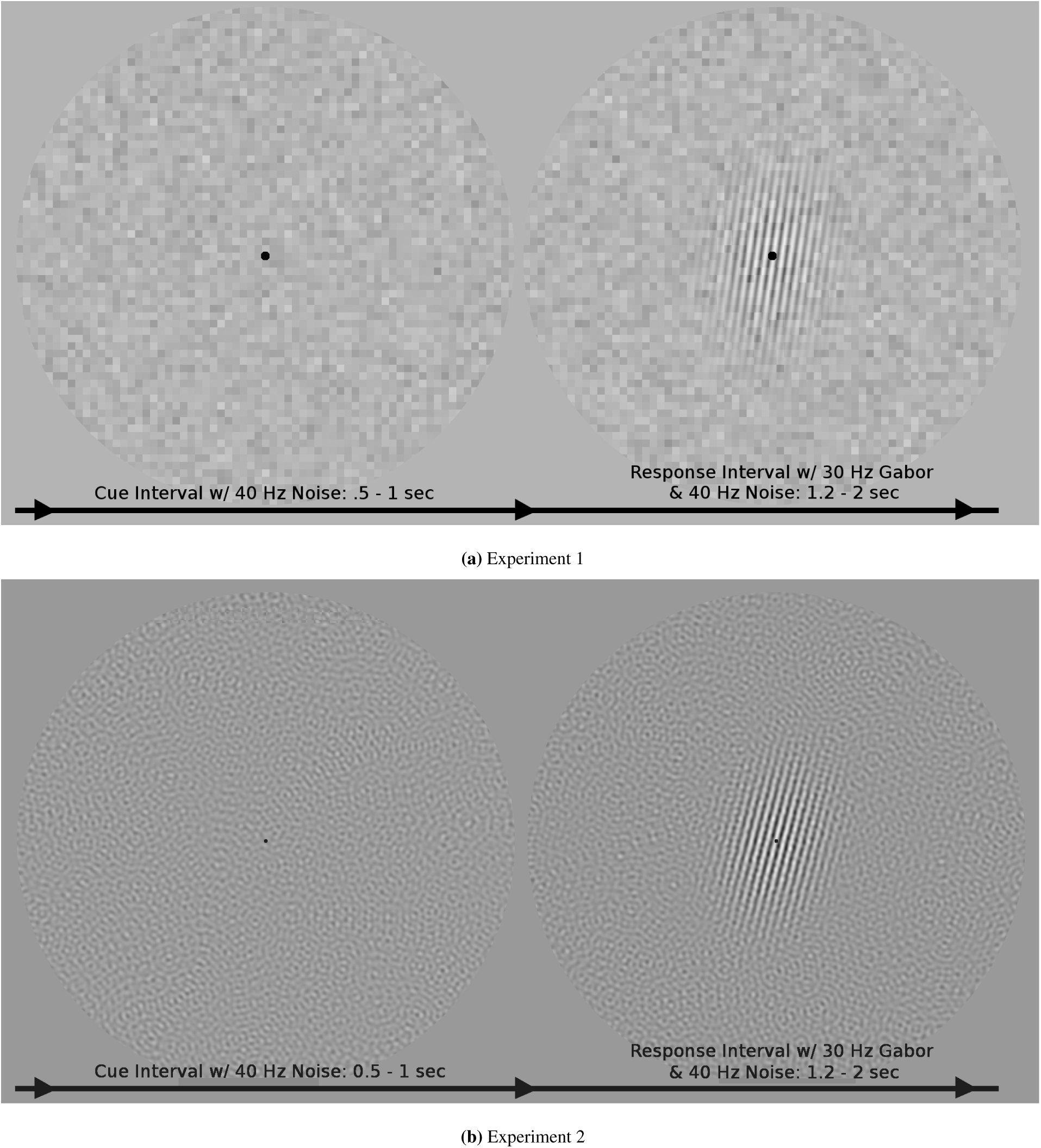
Example stimuli of the *cue* and *response* intervals of medium noise conditions from Experiments 1 (top) and 2 (bottom). During the response interval, participants decided which spatial-frequency-target each Gabor represented, pressing a button with their left hand for a low spatial frequency target (2.4 cycles per degree visual angle, cpd) and pressing a button with their right hand for a high spatial frequency target (2.6 cpd). N200 waveforms were calculated time-locked to the onset of the Gabor stimulus during the response intervals. In both experiments a paradigm was used in which the visual noise changed at 40 Hz and the Gabor signal flickered at 30 Hz to evoke 40 Hz and 30 Hz responses in electrocortical activity that track attention to the noise and signal stimuli. While the attention analysis is beyond the scope of this paper, a similar analysis is presented by Nunez et al. (2015).

Participants performed perceptual decision making tasks in a dark room on a 61 cm LED monitor maintaining a distance of approximately 57 cm from retina to display. The monitor resolution was set as 1920 *\** 1280 pixels with a 120 Hz refresh rate. The participants were told to remain fixated on a small fixation spot while performing the task.

Gabors are sinusoidal grating patterns with a Gaussian falloff of contrast that produces maximal firing in certain neurons within the primary visual cortex (Webster et al., 1985). Participants in both experiments were tasked with identifying the spatial frequency (either “high” or “low” spatial frequency) of large Gabors (approximately 10 cm and 10 degrees visual angle in diameter) that were randomly rotated and had a random phase on each trial (uniformly distributed draws across both rotation angle and phase). In Experiment 2, the high and low spatial frequencies of the target Gabors were 2.4 and 2.6 cycles per degree visual angle (cpd) respectively, while in Experiment 1, the targets were randomly drawn either from one of two shifted beta distributions, ℬ*eta*(13.5, 1.5) distribution plus 1.5 cpd or a ℬ*eta*(1.5, 13.5) distribution plus 2.5 cpd. These two target shifted-beta distributions had means 2.4 and 2.6 cpd respectively and were chosen so that there would exist small variability in the target difficulty (i.e. distance from 2.5 cpd) with two completely separable distributions. Both experiments contained three conditions of high, medium, and low visual noise contrast. Throughout each trial, samples of visual noise were displayed both before, and concurrent with the targets. A new sample was presented at regular intervals (25 ms, every 3rd video frame). In Experiment 1, a checkerboard noise pattern was displayed, while in Experiment 2, the noise was spatially bandpass filtered to include only masking noise centered around 2 and 3 cpd (i.e. the noise was a mixture of two normal distributions centered at 2 and 3 cpd with .1 cpd standard deviations), equally masking both the high and low spatial frequency targets.

The time course of each trial was as follows: participants were asked to fixate on a small fixation spot in the center of the screen throughout the experiment, visual noise changing at 40 Hz was displayed a variable length of time between 500 ms to 1000 ms during the *cue* interval, then the Gabor signal stimulus was flickered at 30 Hz embedded in the noise stimuli for 1200 ms to 2000 ms during the *response* interval. During the response interval, participants were asked to respond as accurately as possible in the time allowed with a button box, using their left hand to respond for low spatial frequency targets and their right hand for high spatial frequency targets. Auditory feedback was given after the response interval to indicate trial accuracy; this feedback was included in order to encourage participant vigilance. An entire session of task performance and EEG collection took participants approximately 1 hour with breaks. Each session produced 8 blocks of 60 trials each for a total of 480 trials, with trials at the three levels of noise contrast intermixed within each block.

### EEG recording

EEG was collected using Electrical Geodesic, Inc.’s 128 channel Geodesic sensor net and a Net Amps 200 series amplifier. Electrical activity from the scalp was recorded at a sampling rate of 1000 samples per second with a vertex reference and hardware bandpass filtered to either 1 to 50 Hz (EEG data from Experiment 1) or 1 to 100 Hz (EEG data from Experiment 2). The hardware band pass filter in Experiment 2 was chosen purposely to maintain high frequencies so that broadband noise was submitted to an Independent Component Analysis (ICA) in order to aid artifact correction. In data from both experiments, visual inspection of the raw time series and resulting ICA components was used to remove obvious artifact due to ocular movements, electrical artifact, electrode movement artifact, and some muscle (EMG) activity (see Makeig et al., 1996; Nunez et al., 2016, for further details), using the MATLAB repository artscreenEEG (Nunez et al., 2017). Both data sets ultimately yielded artifact-resistant EEG data by first removing a small amount of artifactual time segments and then removing some linear electrode mixtures given by clear artifactual Independent Components. The EEG data was then common-averaged referenced and submitted to further processing as discussed below.

### Estimation of N200 waveforms

N200 waveforms, calculated from the EEG signals following the onset of the Gabor patch during the *response* interval were obtained using five processing steps: 1) software bandpass filtering the EEG data from 1 to 10 Hz (forward-backward Butterworth IIR filter with stopbands at 0.25 and 20 Hz with 1 dB attenuation in the passband and 10 dB attenuation in the stopband) obtaining cleaner estimates of the early evoked potentials—note that slower, later evoked potentials were purposefully removed by using a 1 Hz highpass, see Acunzo et al. (2012), 2) subtracting the average baseline potential (i.e. the average amplitude in a 100 ms window before the stimulus), 3) averaging across trials to obtain a traditional ERP estimate at 128 electrodes for each participant (excluding those electrodes marked as artifact), 4) taking a singular value decomposition (SVD) of the ERPs for each experimental condition and EEG session, and then 5) using the first SVD component (the component that explained the most variance) as a channel weighting function in order to obtain a less-noisy estimate of the N200 waveforms for each experimental condition and EEG session as well as to obtain single-trial estimates (discussed below).

The goal of the singular value decomposition (SVD) was to improve the signal-to-noise ratio of both traditional ERP estimates and single-trial estimates by estimating an optimal spatial filter to detect the N200 (Kayser and Tenke, 2003; Parra et al., 2005). SVD is the algorithm used by most principal component analysis (PCA) algorithms and produces non-stochastic, deterministic results. N200 waveforms were obtained by finding the first principal component of the matrix of ERPs of 1000 ms post-stimulus and 100 ms pre-stimulus (samples *T* by channel *C* data, see Methods in Nunez et al., 2017). The first principal component consists of both a weight vector that produces a time series of the trial-averaged ERP and an associated vector of channel weights representing the spatial distribution of that component. Traditionally calculated ERPs (Luck, 2012) at specific parietal electrodes were also compared to the SVD estimated ERP and exhibited no difference in latency.

Single-trial estimates of the SVD-estimated ERP are then obtained by applying the weight vector as a spatial filter (i.e. weight summing over the vector of weights as show by **Figure 2**). It should be noted that the *first* component of the SVD method yielded the N200 waveform in these experiments because 1) the neural response to the visual stimulus was large in the average ERP and 2) other waveform components of the ERP (such as the P300 response) were removed using the 1 Hz highpass. Had the experimental conditions and filtering parameters been different, the N200 waveform may have been placed in a SVD component that explained less of the variance.

**Figure 2.**
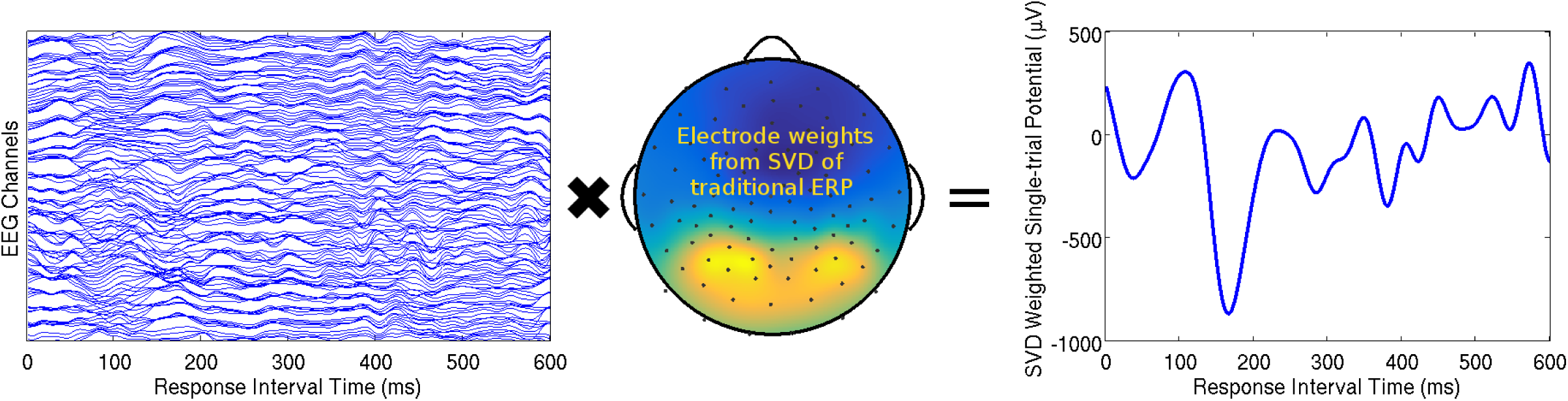
A visual representation of the singular value decomposition (SVD) method for finding single-trial estimates of evoked responses in EEG. The EEG presented here is time-locked to the signal onset during the response interval, such that the single-trial N200 waveform encoded the response to the signal onset. A single trial of EEG from one participant (Left) can be thought of as a time by channel (*T ×C*) matrix. SVD weights **v** (*C ×* 1) are obtained from the ERP response (i.e. trial-averaged EEG; *T ×C*) and can be plotted on a cartoon representation of the human scalp with intermediate interpolated values (Middle). This specific trial’s N200 waveform (Right) was obtained by multiplying the time series data from each channel on this trial by the associated weight in vector **v** and then summing across all weighted channels. Weight averaging across channels is a common method to obtain larger signal-to-noise ratios in EEG (e.g. cleaner, more task-relevant neural signals, see Parra et al., 2005).

The window to calculate minima in order to evaluate N200 peak-latency responses was found empirically. Out of the following windows: 101 to 299 ms post-stimulus, 101 to 249 ms post-stimulus, 151 to 224 ms post-stimulus, and 151 to 274 ms post-stimulus, the 151 to 274 ms window post-stimulus was found to capture the N200 waveform well with few false detections. Only 2 estimates out of 147 estimates of trial-averaged N200 peak-latency were detected at the 274 ms boundary and none at the 151 ms boundary. Single-trial N200 peak-latencies that were found with a minimum on the boundary were removed from the analysis. A minimum at either 151 or 274 ms was indicative that single-trial N200 latencies were not well estimated on that trial, as the interval contains a ramp rather than a peak.

Deflection times were also calculated for the trial-averaged N200 waveforms by taking the window of 0 to 274 ms post-stimulus onset in the response interval and finding were the derivative of the ERP in that window first became negative before the N200 peak-latency. This reflects the point at which the signal begins to decrease before the N200 peak-latency. This particular time point has been explored in a previous study of visual encoding by Martin et al. (2010), albeit calculated using the point at which the signal reached two standard deviations below the baseline. Trial-averaged N200 waveforms and deflection time and peak-latency distributions are shown in **Figure 3**.

**Figure 3.**
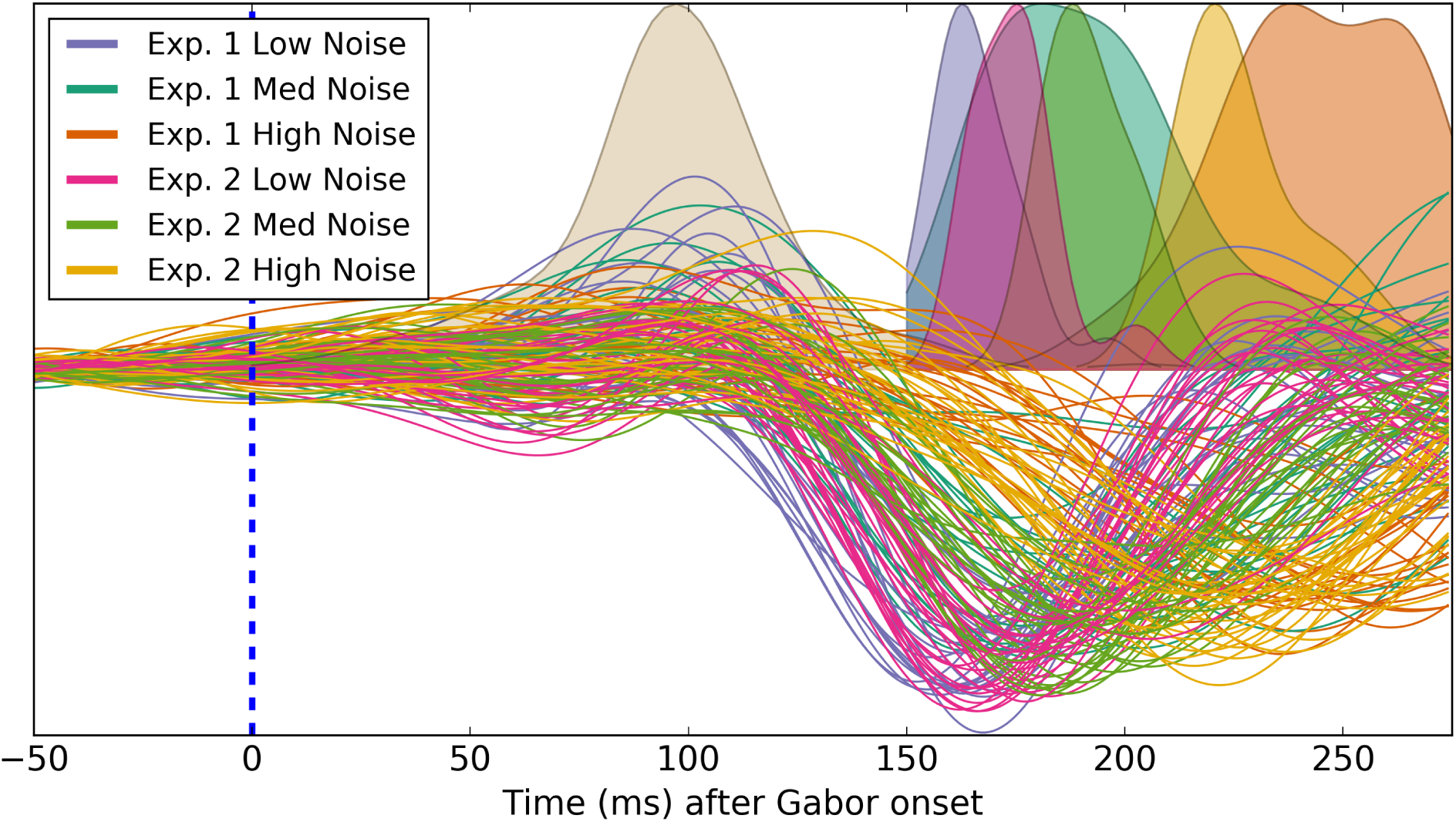
All trial-averaged N200 waveforms across EEG sessions and the six visual noise experimental conditions (*N* = 147). Smoothed histograms for the N200 deflection times (top left distribution) as well as the N200 peak-latency times (all other distributions for each noise condition) are displayed above the zero amplitude midline. There was a significant but weak correlation found between the N200 deflection times and the N200 peak-latency times (*ρ* = .274). N200 peak-latencies were dependent upon the noise condition of the experiment, while N200 deflection times were not found to vary dependent upon the noise condition.

### Cognitive theory of human decision-making

We tested the relationship between N200 peak-latency and VET directly by fitting response time and choice data from the participants to simple drift-diffusion models (DDM). In this theoretical framework, human *visual encoding time* (VET) is the first cognitive processing time after a visual stimulus is displayed, after which the participant accumulates evidence based on the visual input (see **Figure 4**). The *non-decision time* (NDT) τ parameter is the sum of VET and other time-invariant processes across trials not associated with a sequential accumulation of evidence, such as motor execution time (Ratcliff, 1978; Ratcliff and McKoon, 2008). Fitting reaction time and choice behavior with DDM results only in NDT estimates, not direct estimates of the components of NDT, such as VET. We therefore tested whether the linear relationship between N200 peak-latency and NDT had a slope of 1, reflecting the hypothesis that N200 peak-latency is one additive component of NDT.

**Figure 4.**
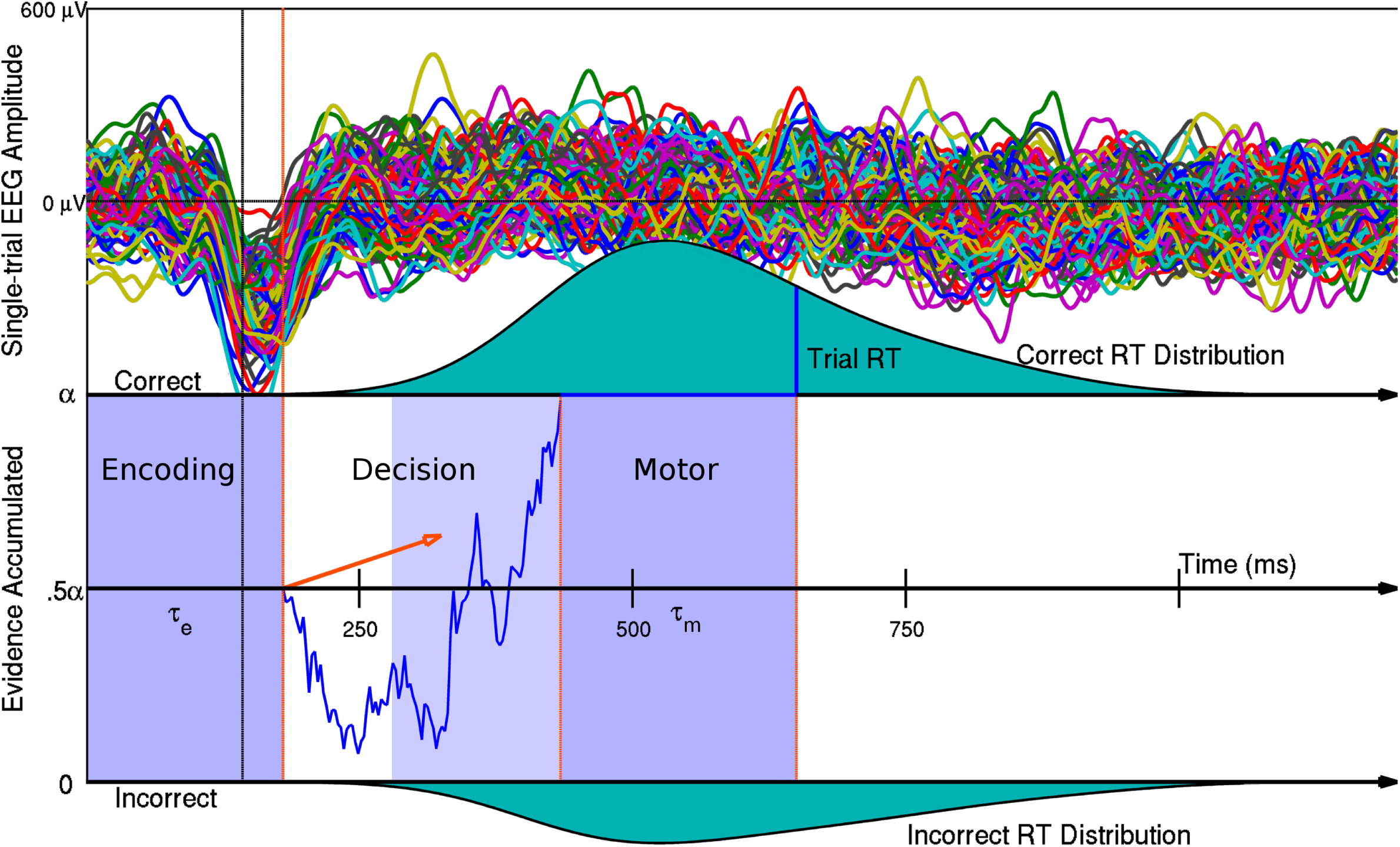
A graphical illustration of a Neural Drift Diffusion model in which the visual encoding time *τ*_*e*_ on single-trials describes the latency of the negative peaks of the N200 waveform on 146 experimental trials. Single-trial observations of the N200 peak-latency are found by using the SVD method (described previously) in one participant and experimental condition. Total NDT τ reflects both visual encoding time (VET) *τ*_*e*_ as well as motor execution time (MET) *τ*_*m*_ (e.g. motor response time of the participant after the decision is made) and can be estimated from response time distributions. Decision time is generated via a Brownian motion evidence accumulation process with evidence accumulation rate δ (drift rate), evidence accumulation noise ς (diffusion coefficient), and evidence threshold (boundary parameter) α (for more information see Ratcliff, 1978; Ratcliff and McKoon, 2008; Nunez et al., 2015, 2017)

We assessed our ability to recover true non-decision time (NDT) estimates from RT percentiles and hierarchical Bayesian model fitting by simulating response time data. Response time data was simulated from drift-diffusion models with various amounts of trial-to-trial variability in NDT, evidence boundary (amount of evidence require to make a decision), and evidence accumulation rate (i.e. drift rate) suggested by Ratcliff (1978) as well as specific amounts of contaminant processes (up to 10% of trials) generating a random choice and reaction time (discussed below). These response times were simulated using DMAT, MATLAB software for simulating and estimating diffusion model parameters (Vandekerckhove and Tuerlinckx, 2008). These simulations revealed that even with the presence of trial-to-trial variability in all cognitive parameters (NDT, evidence boundary, and evidence accumulation rate), means of non-decision time and evidence accumulation rate could still be recovered. In particular, mean NDT across trials was well estimated by 1) 10th percentiles of response time distributions (with about 60 ms of fixed bias in 10th percentiles compared to true mean NDTs) and 2) hierarchical Bayesian fitting of a simple drift-diffusion model (DDM) with between-session variability in the three free parameters (Wabersich and Vandekerckhove, 2014).

It should be noted that mean NDTs are not as well estimated in the presence of a large amount of contaminant trials without explicitly modeling contaminant trials (see **Figure 5**). Contaminant trials are defined as response time observations that are not due to a decision making process and occur due to another random process (e.g. a “contaminant” process unrelated to the theorized cognitive process). Because these response times are not due to a decision process, the associated accuracy on those trials should be about 50% in a two-alternative forced choice task. Recently researchers have developed theory on the notion of “mind-wandering”, which is defined as cognition that departs from that which is useful to the task being completed (e.g. Hawkins et al., 2015; van Vugt et al., 2015; Fox et al., 2015; Lin et al., 2016). We assumed that each reaction time and choice observation was from a mixture model such that some percentage of trials was from a Diffusion Model process and some percentage of trials was from a uniform distribution of reaction times and a binomial distribution of accuracy with parameter θ = .5 (that is, the sample mean of accuracy would be about 50%). Drugowitsch et al. (2012) previously used this method to account for so-called “lapse trials”, although we prefer the term “contaminant” trials in this paper.

**Figure 5.**
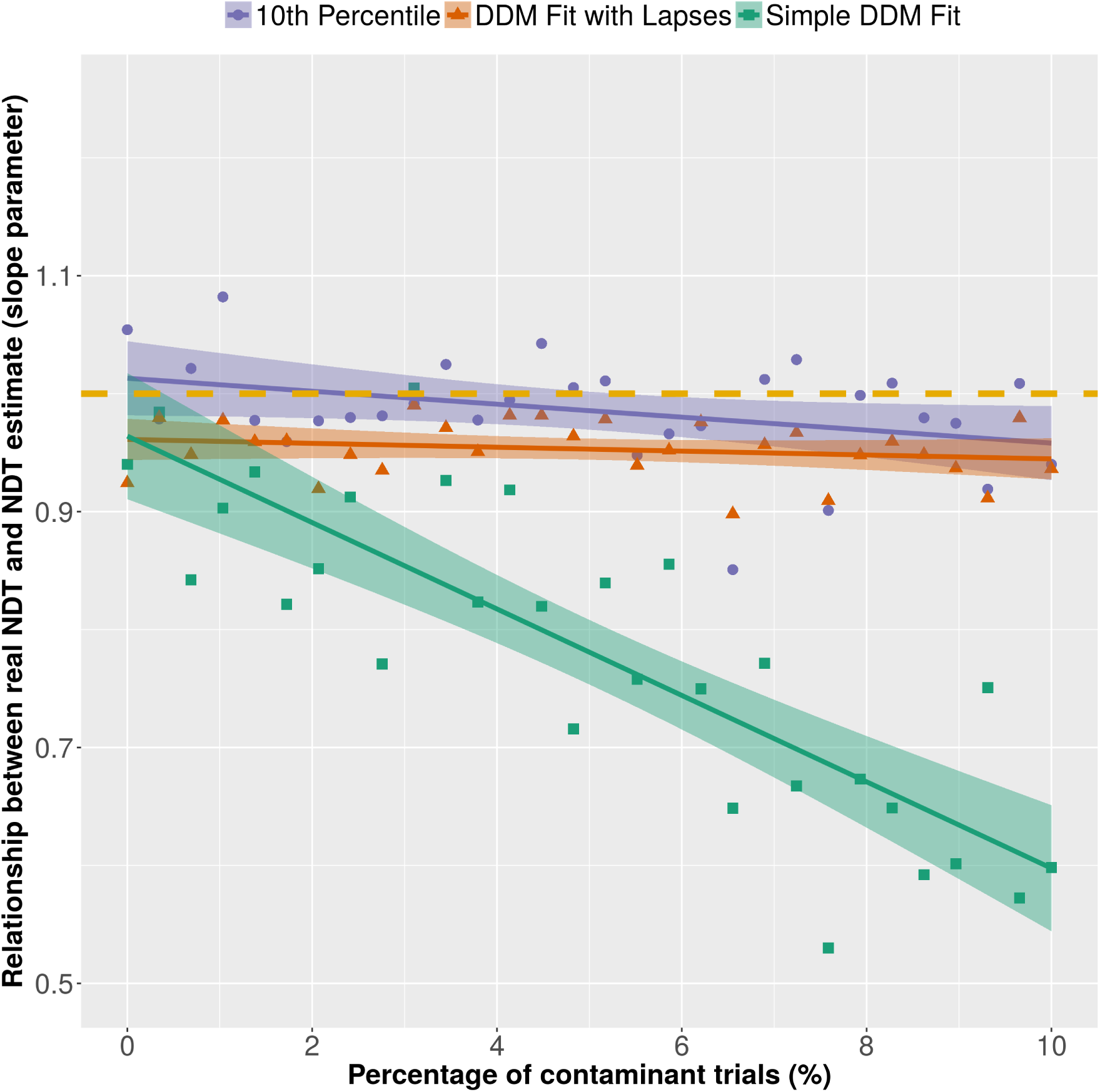
The robustness of non-decision time (NDT) estimation to contaminant processes was evaluated by calculating the slope parameter between true and estimated NDT for 30 simulated participants with 100 trials each, repeated 30 times with increasing amounts of contaminant trials. The dashed gold line represents a slope of 1 and thus would represent recovered NDTs for each participant. Without explicitly modeling contaminant trials, the addition of more trials from contaminant processes resulted in some non-decision time (NDT) estimates being negatively biased towards smaller values, resulting in smaller slope parameters (Green). This simulation also revealed that while 10th response time percentiles were positively biased as estimates of true NDT (e.g. true NDT were approximately 60 ms shorter than the 10th response time percentiles), the slope of 1 between estimated and true NDT relationship could be well estimated by the 10th percentiles (Purple). Explicitly modeling lapse trials (Orange) was shown to be a consistent procedure for finding NDT correlates in EEG data, although the estimated NDT are slightly underestimated.

### N200 waveforms versus response times

To test our hypothesis using a simple method, we performed linear regressions of 1) trial-averaged N200 deflection times and peak-latencies versus 10th response time percentiles across noise conditions and EEG sessions and 2) single-trial N200 peak-latencies versus response times. The fastest response times in each EEG session and noise condition are expected to be the linear sum of *visual encoding time* (VET) and other sequential cognitive processes that are mostly invariant across trials. This is because in any response time process that contains a mix of stochastic evidence accumulation and fixed processes with smaller amounts of variability, NDT (time periods not associated with evidence accumulation during decision making trials, see Voss et al., 2004; Ratcliff and McKoon, 2008) will be reflected in the fastest response times. The fastest response times in each EEG session and noise condition were estimated by the 10th percentile. The 10th percentile was chosen because the smaller percentiles were more likely to include contaminants as discussed above.

Linear regression statistics reported are 95% confidence intervals related to the *p* values and *t* statistics in the Neyman and Pearson (1933) tradition as well as Bayes Factors (BF) which describe the amount of relative evidence (probability in the Bayesian definition) of a model which has a non-zero regression slope over a model which has a regression slope of zero (Jeffreys, 1961; Kass and Raftery, 1995; Rouder and Morey, 2012). Generally Bayes Factors over 3 are considered positive evidence for an effect (i.e. the effect is 3 times more likely under the alternative hypothesis than the null hypothesis) while over 20 is strong evidence (Kass and Raftery, 1995). Adjusted *R*^2^ is also reported that describes the fraction of variance of the dependent variable (10th response time percentiles or single-trial reaction times) explained by the regressor variable (measures of neural processing speed). 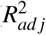 ranges from 0 to 1. These statistics were generated by JASP, a open-source graphical software package for statistical analysis (JASP Team, 2017).

Also reported are Bayes Factors (BF1) of linear relationships between response time and N200 measures that describe the amount of relative evidence of the slope parameter being equal to 1 (e.g. an *x* ms increase in N200 peak-latency corresponds to an *x* ms increase in visual encoding time) over a model which has a largely unknown regression slope. That is, the comparison density and prior probability of slope parameter was *β ∼ 𝒩* (1, 3^2^). BF1 were calculated using the simple Savage-Dickey Density Ratio (Dickey and Lientz, 1970; Wagenmakers et al., 2010) which is the ratio of the height of the posterior distribution over the prior distribution at the tested value. These Bayes Factors were calculated by running simple JAGS code, a program that easily draws samples from marginal posterior distributions using Markov Chain Monte Carlo (MCMC; Plummer, 2003). These Bayes Factors, BF1, are typically much smaller than the Bayes Factors used to test the existence of an effect because less data is needed to prove the existence of an effect than to prove the effect is equal to a specific value (slope equal to 1, in this case).

### Integrated neuro-cognitive model fitting

In order to test our N200-VET hypothesis directly, we fit hierarchical models to behavioral data and assumed linear influences of the N200 peak-latency on cognitive parameters. Thus all linear relationships between N200 peak-latency and NDT in every visual noise condition were estimated in a single-step in a hierarchical Bayesian framework.

Drift-diffusion modeling was applied to response time and accuracy data from Experiments 1 & 2 jointly, containing between session-differences in NDT τ, evidence accumulation rate δ, and speed accuracy trade-off parameter α that were explained differences in N200 waveform peak-latencies *z*. All built models were assumed to be hierarchical, describing intrinsic session *j*, condition *k*, and experiment *e* variability which 1) ensured model fits with small amounts of data (Lee, 2008; Lee and Newell, 2011; Vandekerckhove et al., 2011) and 2) provided the ability to describe data more accurately for observed and unobserved participants in future analyses (Wagenmakers, 2009; Nunez et al., 2017). Bayesian hierarchical drift-diffusion models were fit using Wiener likelihood approximations in JAGS, a program that can easily sample from hierarchical models using Markov Chain Monte Carlo (MCMC) (Plummer, 2003; Wabersich and Vandekerckhove, 2014) using the pyjags Python package (Miąsko, 2017).

Hierarchical Model 1 tested a linear influence of trial-averaged N200 peak-latencies *z*_*jke*_ on each parameter. Hierarchical Model 2 tested the linear influence of single-trial estimated N200 peak-latencies on non-decision time. Notation “*∼*” refers to a probability distribution of data or prior distribution of parameters (either from a normal distribution with notation 𝒩, truncated normal distribution with truncation between *a* and *b* indicated by ∈ (*a, b*), Gamma distribution with notation Γ, or Uniform distribution with notation 𝒰).

Both models assumed that a certain proportion θ of response times (varying between 0% and 100% of trials) came from a contaminant lapse process (discussed above) that produced a random response (Bernoulli distribution with probability parameter *ρ* = .5) and a random (Uniformly distributed) response time between 0 and a large response time (bounded by the maximum observed response time for that EEG session in that visual noise condition). Note that notation *y* ∼ 𝒰 (− max(RT), max(RT)) expresses this information when negative y encode error responses and positive *y* encode encode correct responses as in the package by Wabersich and Vandekerckhove (2014).

In Model 1, wide (uninformative) priors were given to the hierarchical effects *µ*_*i*_ of N200 peak-latency on drift-diffusion model parameters *i*, centered at 1 in order to calculate Bayes Factors (BF1) testing a slope-of-1 relationship using the Savage-Dickey density ratio (Verdinelli and Wasserman, 1995). As a sensitivity analysis, wide priors centered at 0 were used in previous fits of the model, which had inconsequential effects on the posterior distributions (i.e. parameter estimates). The final fit of Model 1 thus had the following embedded linear regression and prior structure:

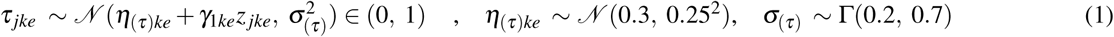

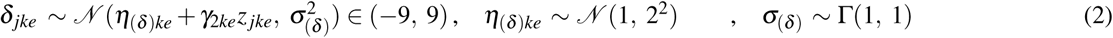

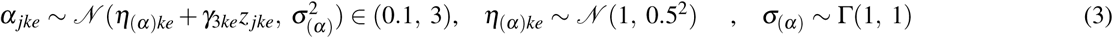

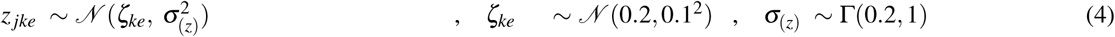

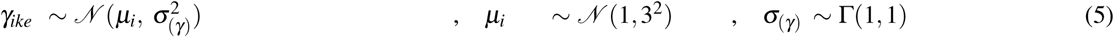

Bayes Factors for each effect parameter *γ* within each visual noise condition and type were calculated using Python and R scripts with the Savage-Dickey density ratio (Verdinelli and Wasserman, 1995) of the posterior density over the prior density at *γ* = 1. These Bayes Factors provide the degree of evidence (defined as a probability ratio in Bayesian statistics) of a model where effect of trial-averaged N200 peak-latency on NDT is 1 versus a model where the effect is unknown (the prior model where *γ ∼ 𝒩* (1, 3^2^)).

The base likelihood that generates observed response time and accuracy data (both represented by vector **y**) is given by the mixture of a DDM and a lapse process described above, with an uninformative prior on the proportion of lapse trials θ for every session *j*, condition *k*, and experiment *e*:

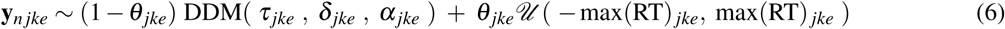

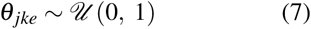

Model 2 was fit to find posterior distributions of decision-model parameters for *single-trial* N200 peak-latency linear effects on single-trial NDT (the sum of VET and motor response time as estimated by a hierarchical Bayesian account of a drift-diffusion model). The model was fit with linear relationships between N200 peak-latencies *x* on single-trials *n* and single-trial non-decision times, a similar framework presented in our previous work (see Nunez et al., 2017). Also included were hierarchical linear effect parameters in order to summarize the linear effects for each noise condition and noise type (i.e. experiment). Model 2 had the following prior structure, a hierarchy very similar to Model 1:

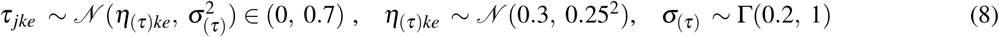

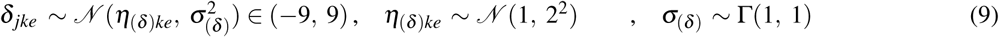

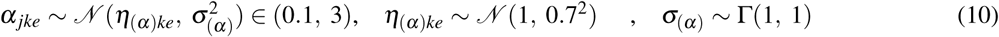

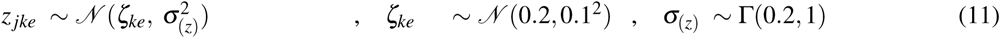

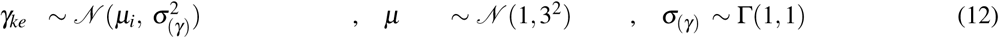

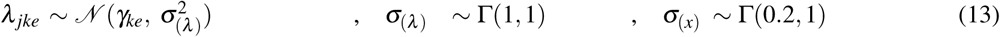

Like Model 1, a lapse process was assumed in Model 2. Model 2 also had a simple normally distributed generative process for the N200 peak-latency *x* on trial *n* truncated between .151 and .274 seconds (see single-trial N200 peak-latency estimation discussion above). Model 2 thus had the following embedded linear regression structure on the likelihood level (where variable **y** refers to the choice and response time on trial *n* from a drift-diffusion model; DDM):

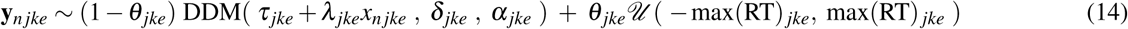

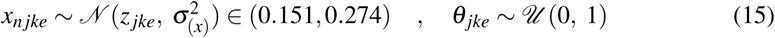

### Location of the N200 waveforms

Spline-interpolation on the scalp surface has been shown to be accurate for high-density EEG (approximately *>* 64 electrodes; Nunez and Srinivasan, 2006). To view the scalp representation of the observed N200 waveforms, we used spline-interpolation in each EEG session to estimate the potential between electrode contacts and to generate interpolated data for artifactual electrode contacts. This method yielded the best representation of the N200 peak-latency on the surface of an MNI human scalp (e.g. see the Top panel of **Figure 9**).

**Figure 9.**
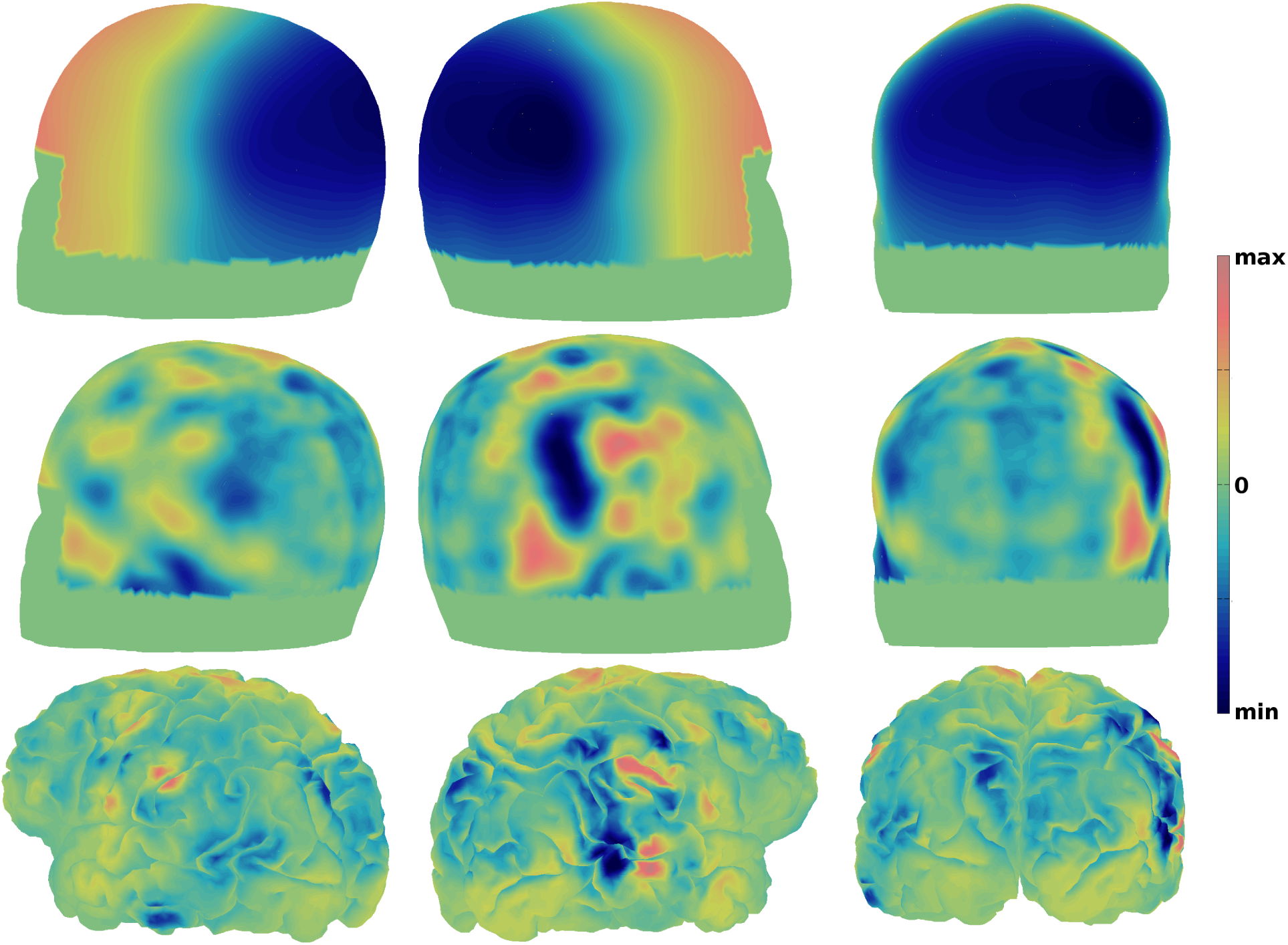
Left and right sagittal and posterior views of localized trial-averaged N200 waveforms averaged across all EEG recordings and noise conditions (*N* = 147). (Top) MNI scalp potential topographies (averaged projections of the first SVD components at the N200 peak-latency) were generated using spline-interpolation between electrodes. (Middle) Current Source Density MNI scalp topographies were generated with a Laplacian transform of the aforementioned data (Nunez and Pilgreen, 1991; Deng et al., 2012; Kayser and Tenke, 2015). (Bottom) The “Representative” cortical maps were obtained by projecting the MNI-scalp spline Laplacians onto one participant’s anatomical fMRI image via Tikhonov (L2) regularization, maintaining similar distributions of activity of the surface Laplacians on the cortical surface. Blue and orange regions correspond to areas that produce negative and positive potentials observed on the scalp respectively.

EEG localization to a three-dimensional representation of the brain is an inexact process that is unsolvable without additional assumptions (Nunez and Srinivasan, 2006). However the surface spline Laplacian (i.e. “Current Source Density”, CSD) on a curved two-dimensional space does not require additional assumptions, has been shown to match closely to simulated cortical-surface activity using forward models (i.e the mapping of cortical activity to scalp potentials), are unaffected by reference electrode choices, and have shown consistent results when used with real EEG data (Nunez and Srinivasan, 2006; Kayser and Tenke, 2015). In addition, unlike 3D solutions, projections to the dura mater surface with CSD are theoretically solvable, and have been used with success in past studies (see Nunez et al., 1994, for an example). Topographic maps of the surface spline Laplacian projection of the SVD-localized, average N200 waveform across EEG sessions and noise conditions are shown in the Middle panel of **Figure 9**. It should be noted that the minimum of the average true-ERP across all observations (*N* = 147) yielded a very similar Laplacian map; although we expect the Laplacian of the true-ERP to yield surface maps that partially reflect some unrelated cognitive processes

Repeating a similar “source-localization” technique found in Nunez et al. (2017), we localized the curved 2D surface spline Laplacians (Deng et al., 2012) to a folded cortical surface, which resulted with most amplitude existing on the gyral crowns (i.e. the “smooth” cortex). The surface spline Laplacians were projected onto a cortical shape generated by an anatomical MRI of a single individual. This individual’s brain image had cortical folds that better localized particular regions than the MNI brain shape. Each participant’s N200 projection was performed using Tikhonov (L2) regularization to inverse the Finite Element (FE; Pommier and Renard, 2005) forward model based on the MNI 151 average head, maintaining similar spatial distributions of activity of the surface Laplacians to the cortical surface. Cortical topographic maps of the average N200 waveform across EEG sessions and noise conditions are shown in the Bottom panel of **Figure 9**. It should be noted that these projections to the cortical surface are only “Representative” sources in the framework by Nunez et al. (2018) and cannot be “Genuine” or “Equivalent” sources.

In the representative cortical maps each brain vertex was labeled using the Destrieux cortical atlas (Fischl et al., 2004). The locations described below were found in at least 70% of localizations across EEG sessions and experimental conditions using an empirically found cutoff (estimated at −10 units microamperes per mm^2^). However we should note that the localization procedure must have some errors due to between-participant variance in both cortical and cranial shape and between-participant variance in tissue properties. These sources of variance could not easily be accounted for, and remains a source of error in this (and all) cortical projection techniques.

### Code Accessibility

Pre-calculated EEG measures, raw behavioral data, MATLAB stimulus code, and MATLAB, Python, R, and JAGS analysis code are available on https://osf.io/jg9u3/ and in the following repository https://github.com/mdnunez/encodingN200 (as of March 2018).

## Results

### Data summaries: Longer N200 peak-latencies with more difficult visual noise conditions

Data summaries of N200 deflection times, trial-averaged N200 peak-latencies, 10th response time percentiles (shown to be a stable but biased estimate of NDT in **Figure 5**), median response times across trials, and mean accuracy across trials are given in **Table 1**, such that the total sample size of each measure was *N* = 147. A significant but weak correlation was found between the found N200 deflection times and the N200 peak-latencies (*ρ* = .274, *p* < .001). A significant correlation was found between the N200 peak-latencies and 10th response time percentiles (a less specific test of our main hypothesis; *ρ* = .387, *p* < .001). No correlation was found between the N200 deflection times and 10th response time percentiles (*ρ* = −0.086, *p* = .30). From the data summaries in **Table 1** it is clear that N200 peak-latencies, 10th response time percentiles, and median reaction times generally become longer in higher-contrast noise conditions, an expected effect of visual noise condition on VET (due to more difficult figure-ground segregation). This was not true between the Low and Medium noise conditions in Experiment 1, although it should be noted that there was not much behavioral difference between Low and Medium noise contrast in Experiment 1 in response time nor accuracy. Decreasing accuracies with higher noise contrast are an expected effect of visual noise contrast on decision-making. The effects of visual noise condition on VET versus decision-making time (both components of response time) were separated by the integrated neuro-cognitive modeling results presented below.

**Table 1.** Summary statistics for each data type used in this study across EEG sessions and visual noise conditions for a total sample size of 147. Experiment 1 had a sample size of 63 (21 unique EEG sessions per visual noise condition). Experiment 2 had a sample size of 84 (28 unique EEG sessions per visual noise condition). N200 peak-latencies, 10th percentile response times, and median response times become longer in more difficult (lower accuracy) noise conditions. This is not true of N200 deflection times in Experiment 2. VET is expected to become longer when segregation of the visual target is more difficult in conditions of high noise contrast.

|  |  | N200 | N200 | 10<sup>th</sup> Percentile | Median | Mean |
| --- | --- | --- | --- | --- | --- | --- |
|  |  | Deflection times | Peak-latencies | Response times | Response times | Accuracy |
| All observations | Overall Mean | 95.5 ms | 199.6 ms | 575.0 ms | 852.8 ms | 78.76% |
|  | 5<sup>th</sup> Percentile | 48.3 ms | 162.0 ms | 443.6 ms | 629.5 ms | 58.36% |
|  | Median | 97.0 ms | 192.0 ms | 562.4 ms | 845.8 ms | 78.76% |
|  | 95<sup>th</sup> Percentile | 136.5 ms | 259.8 ms | 733.6 ms | 1139.5 ms | 95.32% |
| Experiment 1 | Low Noise Mean | 81.9 ms | 166.3 ms | 549.1 ms | 833.0 ms | 72.25% |
|  | Medium Noise Mean | 94.0 ms | 193.0 ms | 541.9 ms | 821.3 ms | 70.14% |
|  | High Noise Mean | 108.2 ms | 244.5 ms | 613.0 ms | 896.8 ms | 61.77% |
| Experiment 2 | Low Noise Mean | 100.1 ms | 175.0 ms | 545.3 ms | 809.1 ms | 90.07% |
|  | Medium Noise Mean | 93.6 ms | 192.2 ms | 565.3 ms | 829.7 ms | 89.59% |
|  | High Noise Mean | 94.5 ms | 228.0 ms | 629.9 ms | 925.3 ms | 80.72% |

There were 14 unique participants across both experiments (with two participants participating in both experiments). In order to explore true individual differences, observations of the N200 deflection time, N200 peak-latency, 10th response time percentiles, median response times, and accuracies were averaged across the two experiments and all conditions for each participant. The range of these measures are given in **Table 2**. These measures varied across participants with a 18 ms standard deviation of N200 deflection time, 12 ms standard deviation of N200 peak-latency, 68 ms standard deviation of 10th response time percentiles, 125 ms standard deviation of median response times, and 9% standard deviation of accuracy across participants. These results suggest individual differences in both VET and decision-making time (e.g. response time related to a accumulation of evidence). Some of these individual differences account for the variation observed in N200 deflection times and N200 peak-latency distributions presented in **Figure 3**. It should be noted that four of the participants were considered trained on the task, with two participants completing 9 sessions (both experiments) and two other participants completing 7 sessions of the second experiment. While not the focus of this particular study, future work should seek to discover if individual differences in VET are explained by individual differences in N200 peak-latencies (a hypothesis expected from the results of the research presented here). Assuming our hypothesis is true, individuals are expected to have larger differences in their decision-making time than VET.

**Table 2.** Summary statistics for each data type averaged within each of the 14 unique participants. All measures varied across participants suggesting individual differences in VET and other decision-related cognitive processes.

| Data summary across unique subjects ($N = 14$) | | | | | |
| --- | --- | --- | --- | --- | --- |
|  | N200 | N200 | 10<sup>th</sup> Percentile | Median | Mean |
|  | Deflection times | Peak-latencies | Response times | Response times | Accuracy |
| Minimum | 57.7 ms | 183.2 ms | 482.3 ms | 649.1 ms | 62.67% |
| Mean | 91.7 ms | 199.5 ms | 567.3 ms | 839.7 ms | 72.33% |
| Maximum | 129.2 ms | 225.3 ms | 676.2 ms | 1070.8 ms | 90.14% |

### Simple results: Linear regressions suggest N200 peak-latencies shift reaction times

Testing the hypothesis that trial-averaged N200 peak-latency predicts session-level visual encoding time (VET), linear models between N200 peak-latencies and 10th response time percentiles (both correct and incorrect response times combined) were fit. In evidence accumulation models of quick decision making, the *shifts* of response time (RT) distributions (and thus the small RT percentiles) should correlate with NDT (i.e. VET + motor execution time) while the shape of RT distributions (and thus the medium and larger RT percentiles) should correlate both with decision-making processes. Thus linear regressions can be explored without fitting decision-making models to data directly. Using simple linear regression, a slope close to 1 was found between trial-averaged N200 latencies and 10th RT percentiles as shown in the top right of **Figure 6** with a regression coefficient of *β*_1_ = 1.14 (*N* = 147, 95% confidence interval: [0.65, 1.63], *p* < .001, *t* = 4.56, BF = 1.58 *\** 10^3^, BF1 = 10.26). Only 12% of the variance in 10th RT percentiles were explained by the variance across trial-averaged N200 latencies 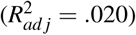. This latter finding might be expected if other trial-invariant mechanisms besides visual encoding time (VET) contribute to the variance in NDT, such as motor execution time. While other regressions between trial-averaged N200 peak-latencies and 30th, 50th, 70th, and 90th RT percentiles and RT means were significant with 95% confidence intervals that included 1, these regressions explained less than 13% in those RT measures. 1st RT percentiles were found to be explained by N200 peak-latencies only when 350 millisecond cutoff values were used to remove some contaminant trials, otherwise the regression was not significant.

**Figure 6.**
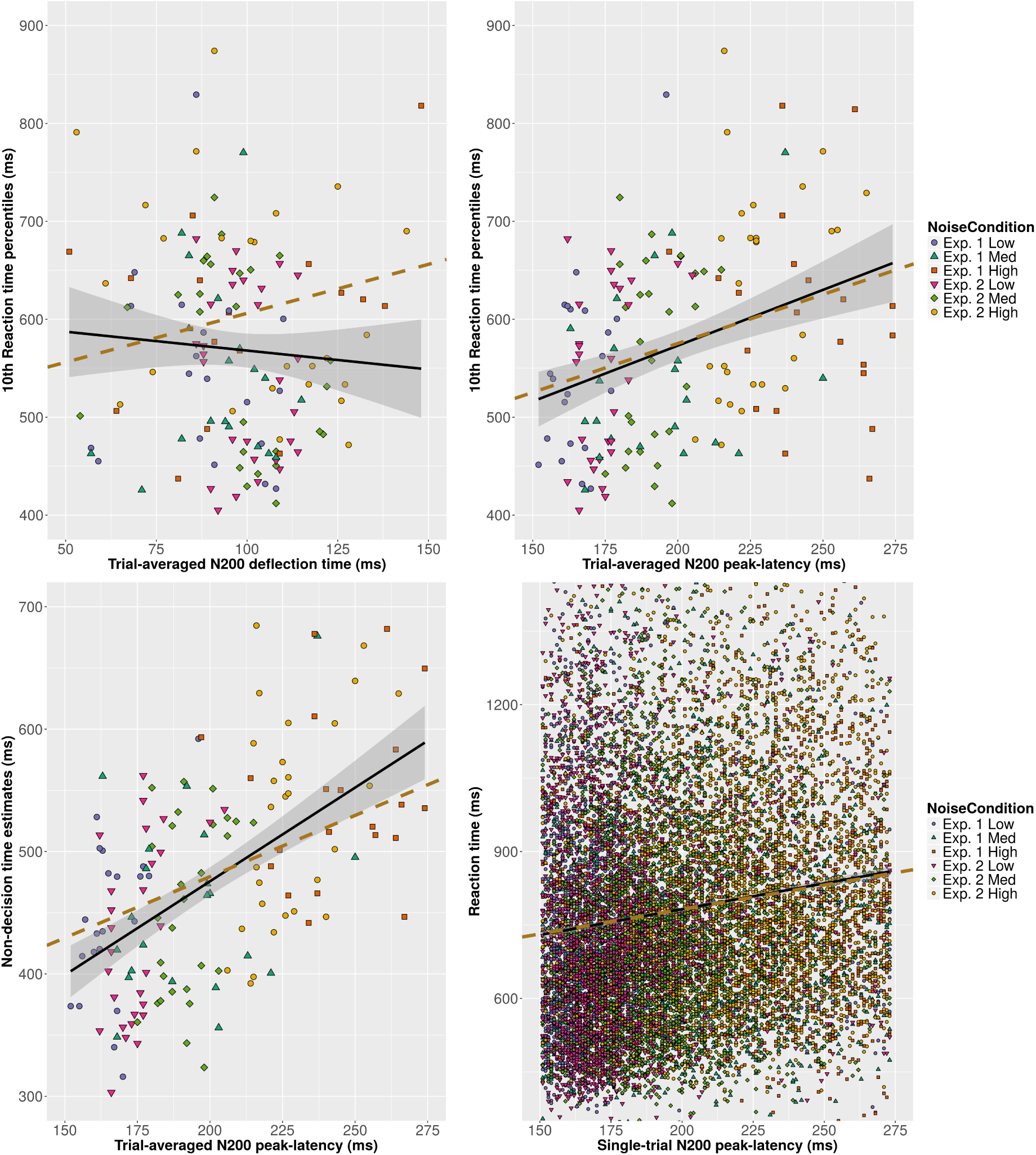
Top Left: A scatter plot of N200 deflection times and 10th response time percentiles (an estimate of non-decision time; NDT) Observations were generated per noise condition and per EEG collection session (*N* = 147). Top Right: A scatter plot of N200 peak-latencies and 10th response time percentiles. Bottom Left: A scatter plot of N200 peak-latencies and NDT estimated using the medians of posterior distributions from Model 1. Bottom Right: A scatter plot of single-trial N200 latencies versus single-trial response times. Observations were generated per trial (*N* = 13,462). Best-fit simple linear regression lines are shown in black with 95% confidence intervals for the intercept and slope parameters shown in gray. Overlaid on the linear regression lines are gold dashed lines representing the hypothesis with a slope of 1.

The relationship is also expected to be observed on single-trials, albeit with more variance around the regression line. Single-trial estimates of N200 latencies were compared to response times across all data points. A slope near 1 was found between single-trial N200 peak-latencies and response times with a tight confidence interval as shown in the bottom right portion of **Figure 6** with a regression coefficient of *β*_1_ = 1.05 (*N* = 13,462, 95% confidence interval: [0.93, 1.18], *p* < .001, *t* = 16.65, BF = 6.30 *\** 10^57^, BF1 = 31.09). However only about 2% of the variance of raw single-trial response times were explained 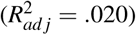. This small amount of variance explained is expected since evidence accumulation should contribute to most of the variance in response times on single trials rather than visual encoding time. This was explored directly with integrated cognitive modeling procedure.

No relationship was found between N200 deflection times and 10th RT percentiles (*N* = 147, *p* = .251, *t* = −1.15, BF = 0.33, BF1 < .001). The linear regression is shown in the top left of **Figure 6**. In fact, the Bayes Factor BF1 displayed evidence strongly in favor of *β*_1_ ≠ 1 (1*/*BF1 = 1.05 *\** 10^4^). This suggests that while N200 deflection times may reflect an early visual processing stage, variance in N200 deflection times across noise conditions and EEG sessions does not correspond with variance in response time distribution shifts across noise conditions and EEG sessions.

### Modeling results: Direct evidence that N200 peak-latencies track visual encoding time

Each model was fit using JAGS with six Markov Chain Monte Carlo (MCMC) chains of 52,000 samples each run in parallel with 2,000 burn-in samples and a thinning parameter of 10 resulting in 5,000 posterior samples in each chain. The posterior samples from each chain were combined to form one posterior sample of 30,000 samples for each parameter. Model convergence was evaluated based on the Gelman-Rubin statistic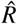, which compares the estimated between-chains and within-chain variances for each model parameter (Gelman and Rubin, 1992). Using this statistic both models were judged to have converged, with 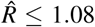 for all parameters in each model (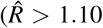 *>* 1.10 for any parameter is usually deemed evidence for a lack of convergence of the entire model fit).

Model 1 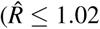 for all parameters) was used to evaluate the effect of trial-averaged N200 peak-latency on NDT across EEG sessions and noise conditions. The mean effect across all conditions of each experiment *µ*_*γ*_ provides a sense of how N200 peak-latency tracks NDT τ in general. The posterior distribution of this parameter was found to be around 1 in each condition and experiment as shown in **Figure 7** and the bottom left portion of **Figure 6**, indicating that an *x* ms increase in N200 peak-latency corresponded to a roughly an *x* ms increase in NDT. The posterior distribution of the overall hierarchical effect of N200 peak-latency on NDT across sessions is plotted at the top of **Figure 7**, indicating that the overall Bayes Factor for a slope-of-1 relationship between N1 latency and NDT is BF1 = 4.62, reflecting moderate evidence (median = 1.39, 95% credible interval: [0.62, 2.26]). However all the posterior distributions are centered above 1 for each unique noise condition. This is expected if N200 peak-latencies reflect VET and estimated NDT are slightly underestimated when including a contaminant process as in Model 1, which is shown by the Orange regression line in **Figure 5**. The effects of N200 peak-latency on two decision-making parameters was also explored in order to disentangle the three cognitive parameters. An undetermined effect of trial-averaged N200 peak-latency on evidence accumulation rate δ was found (median = −1.53 evidence units per second, 95% credible interval: [−5.38, 2.27]). An undetermined effect of trial-averaged N200 peak-latency on boundary separation α was found (median = 0.69 evidence units per second, 95% credible interval: [−0.93, 2.44]), assuming boundary separation is well estimated (which was not shown in simulation, see Methods).

**Figure 7.**
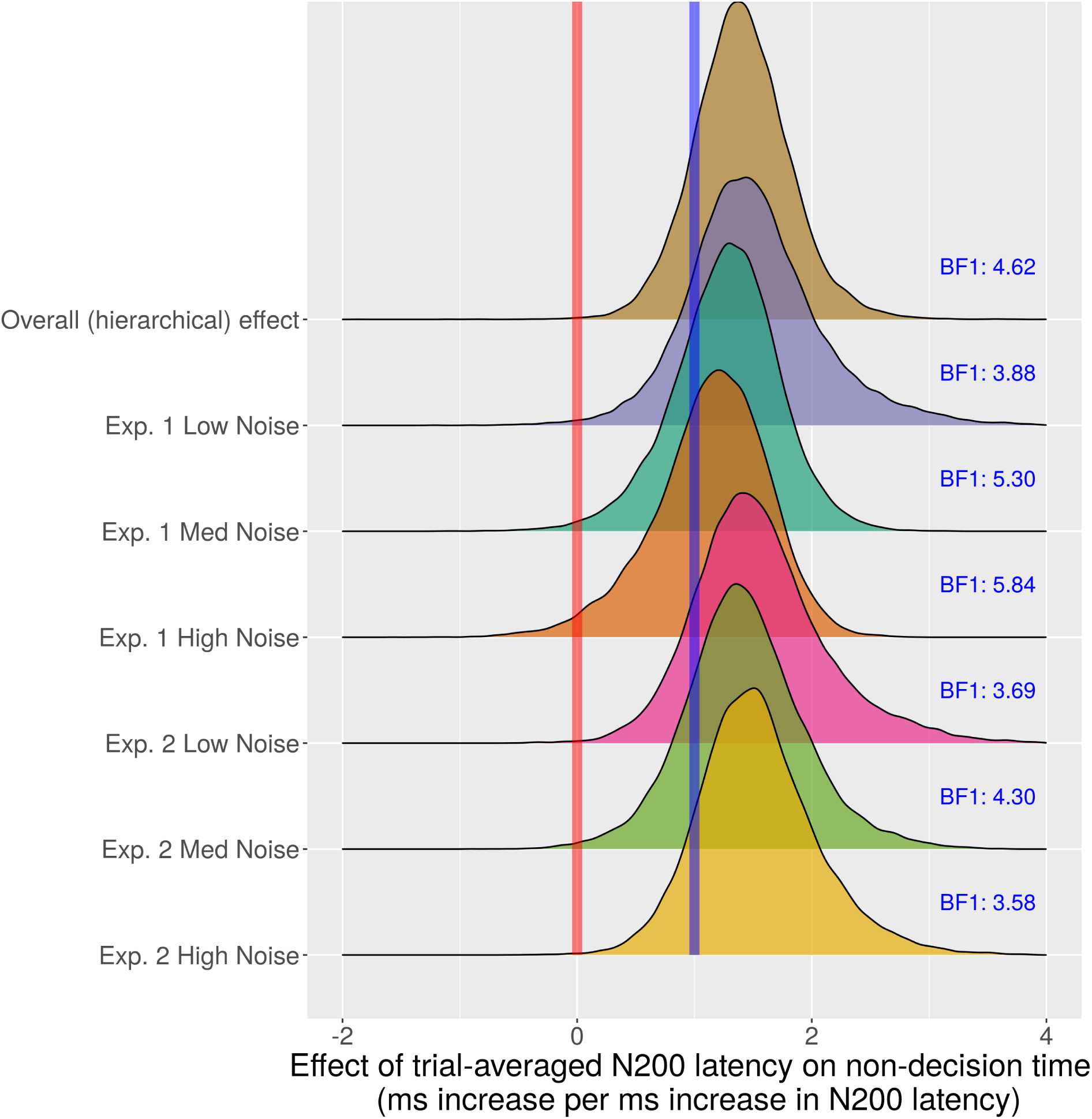
Posterior distributions (Model 1) of effects of N200 peak-latency on non-decision time (NDT). Posterior distributions provide measures of parameter uncertainty in Bayesian model-fitting methods. If the posterior distributions had most of their density around the blue line, this would be strong evidence for the slope-of-1 relationship when the model assumptions are true. Posterior distributions with most of their density around the red line would be strong evidence for the null hypothesis (the parameter equals 0). Some evidence exists for the effects of N200 peak-latency on NDT to have a slope of 1 (*x* millisecond increase in N200 peak-latency corresponds to a *x* millisecond increase in NDT) as indicated by the Bayes Factors calculated with a Savage-Dickey density ratio (Verdinelli and Wasserman, 1995) of the posterior density over the prior distribution at *γ* = 1.

The residual non-decision time *η*_(*τ*)_ posterior distributions yield predictions about the motor execution time (MET), thought to be the second component of NDT that is not VET. Even though NDT changes based on the visual noise condition, these changes should be reflected in the VET (and thus N200 peak-latencies) and not the motor execution time (MET). Thus the residual non-decision time should be similar in each visual noise condition. This is indeed the case, with all residual non-decision time posterior distributions overlapping significantly (see **Figure 8**), reflecting a large probability that the residual NDT is equal in each noise condition. While the median posterior distributions range from 170 to 260 milliseconds, the best estimate of MET is given by the condition with the slope parameter posterior closest to 1, implied by the High Noise contrast condition in Experiment 1. This suggests that the best estimate for MET is 260 milliseconds, which matches estimates for MET we found in another study by Lui et al. (2018).

**Figure 8.**
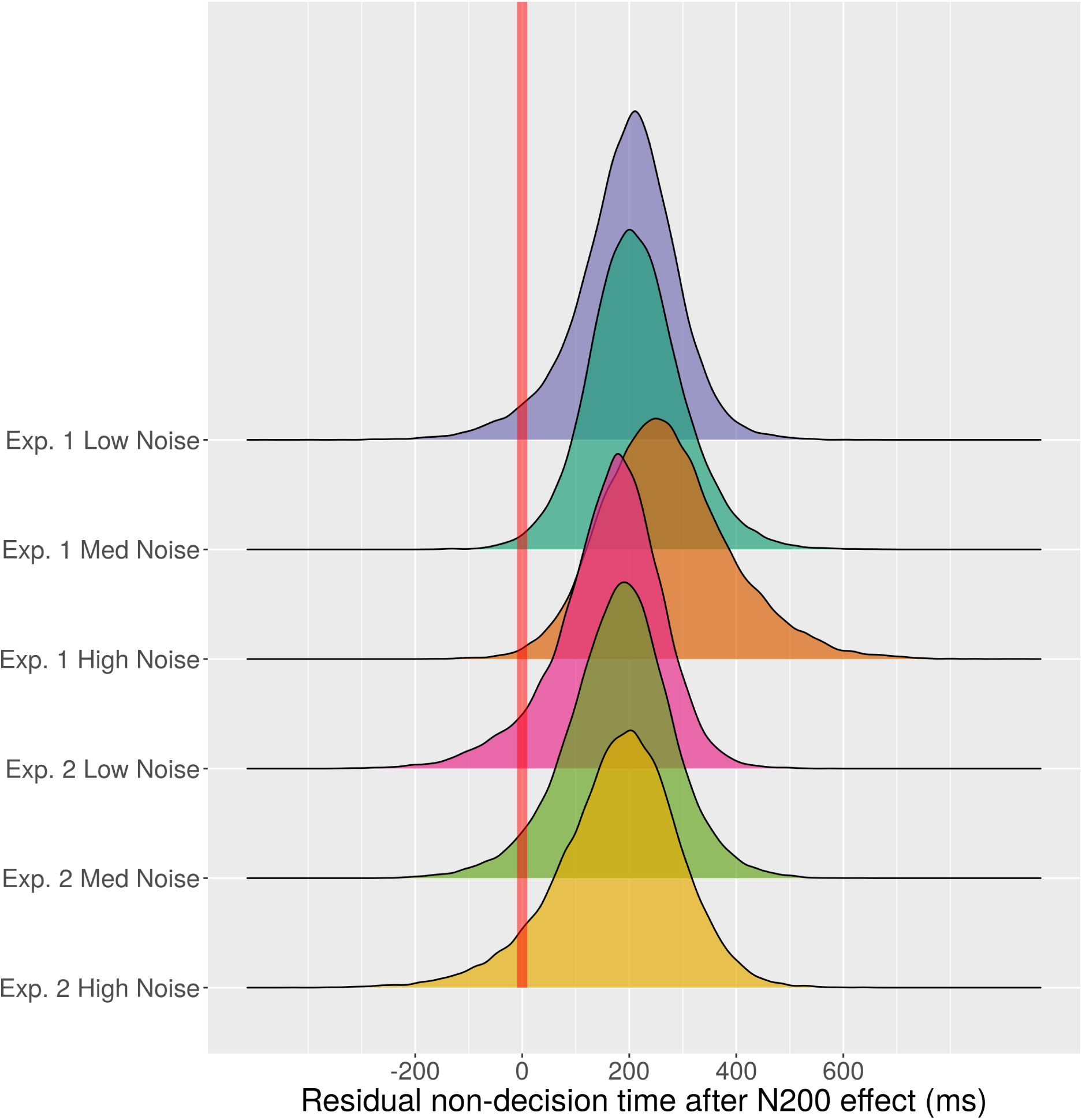
Posterior distributions (Model 1) of residual non-decision time (NDT) after accounting for the effect of N200 peak-latency. The N200 peak-latency explains most of the variance in non-decision time across visual noise conditions. The residual non-decision time is thought to be an approximate measure of motor execution time (MET) and is not expected to depend upon the visual noise condition. As expected, the posterior distributions are similar in each visual noise condition, yielding only minor evidence for a difference in motor execution times between the two experiments.

Model 2 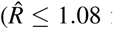 for all parameters) was fit in order to evaluate the effect of single-trial N200 peak-latency on non-decision time for each EEG session. In Model 2, the hierarchical effect parameter *µ*_*γ*_ provides a sense of how *single-trial* N200 peak-latency effects NDT. This parameter suggested that the overall single-trial effect was positive (median = 0.34, 95% credible interval: [0.22, 0.46]). This positive relationship on single-trials between N200 latency and NDT replicates our previous finding in different dataset (Nunez et al., 2017). However no evidence was found that this single-trial effect was equal to 1 (BF1 < .001). That is, there was strong evidence *against* a slope of 1 on single-trials between N200 peak-latency and NDT. Model 2 necessarily does not allow for measurement error in N200 peak-latencies on single trials. Model 2 thus may not accurately capture a true correspondence to VET when the single-trial neural measure contains measurement noise (see Hawkins et al., 2017, for a discussion on noisy neural measures embedded in cognitive models)

### Brain networks that may generate N200 waveforms

N200 peak-latencies are distributed across the cortex and not localized to one particular area. This is reflective of a network that gives rise to the N200 peaks instead of local processing. The distributed network is evident from different focal regions of the average N200 waveform in the spline Laplacian (see Middle portion of **Figure 9**). This is also evidenced by the deviating time course of all 128 electrode locations at the N200 peak latency in the spline Laplacian of each traditional ERP, as shown in one example in **Figure 10**. The spline Laplacian results leads us to reject the hypothesis of a single generator of N200 peak-latencies. This is also true of the localization presented at the Bottom of **Figure 9**, however the sources estimated are only representative (see Nunez et al., 2018) and may not be that precise.

**Figure 10.**
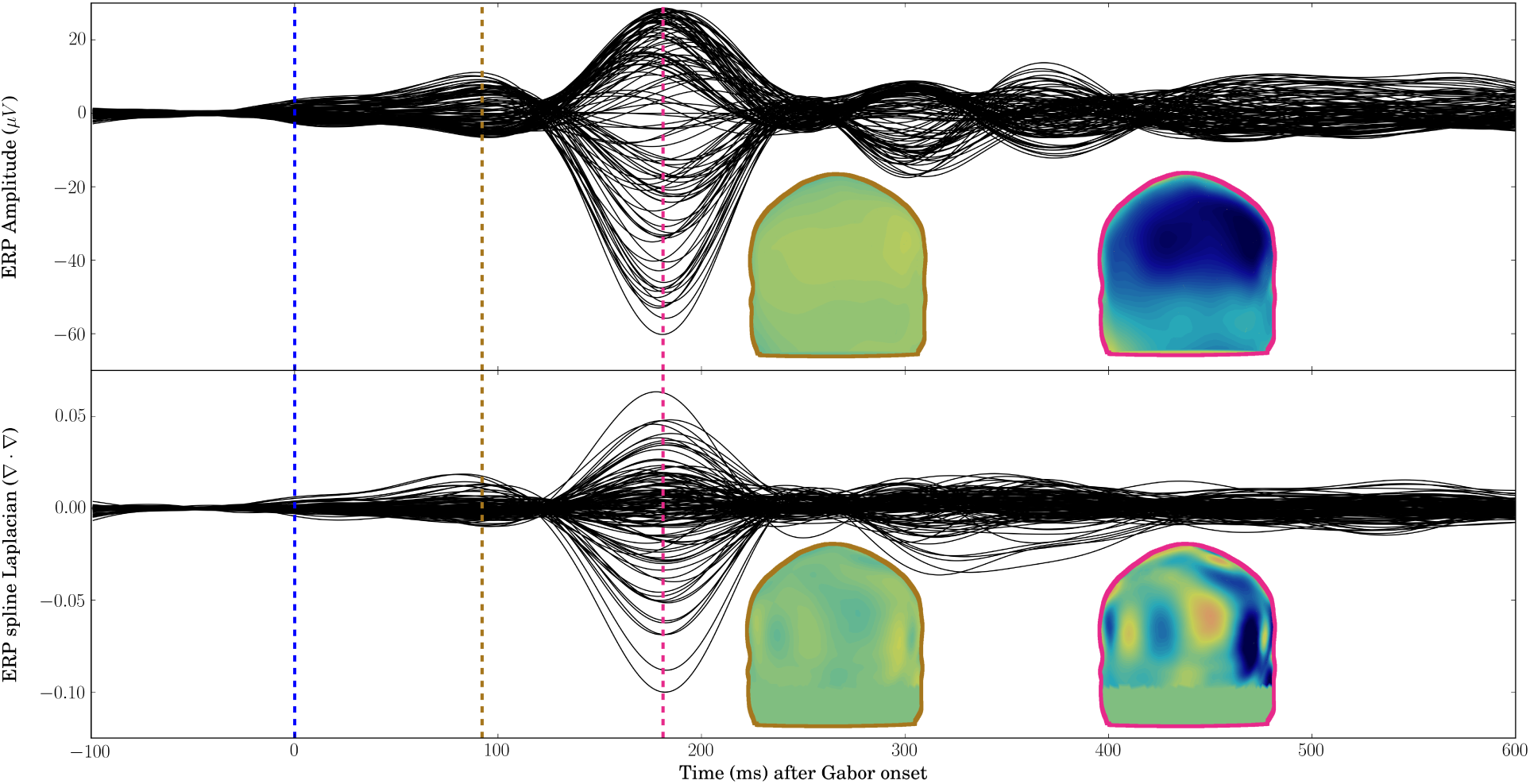
The time course of one indicative traditional ERP (Top) from Experiment 2 in the Medium noise condition for 128 electrode locations filtered from 1 to 10 Hz and spline Laplacian transform of the same ERP (Bottom) at the same locations. ERP potentials and spline Laplacians are shown on posterior views of the MNI scalp at 92 milliseconds (Gold, left) and 181 milliseconds (Pink, right) using spline interpolation between electrode locations. The variability across electrode locations of the spline Laplacian transform at 181 ms is indicative of a distributed network that gives rise to the N200 peak-latency. The topography of average SVD-localized N200 waveforms is shown in the middle portion of **Figure 9**.

While the estimated cortical sources were distributed, the source estimation method described above suggested that N200 waveforms in this study could be found primarily in the following cortical locations (such that the locations listed here were present in at least 70% of the localizations across EEG sessions and noise conditions): the middle temporal gyri (both hemispheres), the postcentral gyri (both hemispheres), and the right angular gyrus. The right middle temporal gyrus was the location of the minimum amplitude (i.e. the greatest magnitude of the N200 peak) in 55.8% of observations across EEG sessions and noise conditions. While all other locations contained the largest negative amplitude in less than 10% of observations. Work by Mishkin et al. (1983) and Lamme and Roelfsema (2000), among others, suggest that the segregation of the target Gabor from visual noise would occur in the temporal “segregation” pathway of the primate cortex at about 150 ms (after a delay due to feedforward and feedback visual network behavior). The possible localization of the N200 waveforms in human extrastriate and temporal locations (especially the right middle temporal gyrus) is further evidence that the N200 peak-latency is the time point at which the target Gabor is “segregated” from visual noise in the human cortex, in order to accumulate visual evidence for a motor response.

## Discussion

Evidence is presented both for and against the N200-VET hypothesis. Evidence against hypotheses are often left out of papers presenting initial results in both the Cognitive Psychology and Cognitive Neuroscience fields, often presented as confirmatory results. Such “censoring” of data can lead to biased statistical analyses (Sterling, 1959; Rosenthal, 1979; Guan and Vandekerckhove, 2016).

### Evidence FOR N200 peak-latencies tracking visual encoding time

Both simple regression analyses yielded evidence that EEG measures of neural processing speed (N200 peak-latencies) track visual encoding times (VET). These regression fits indicate that there was 1) strong evidence for an *x* ms increase in response times when there was an *x* ms increase in single-trial N200 peak-latency and 2) evidence for an *x* ms increase in the 10th percentile of response times, an estimate of cognitive non-decision time (NDT), when there was an *x* ms increase in trial-averaged N200 peak-latencies. Furthermore, moderate evidence was found for a slope of 1 when fitting embedded linear regression of trial-averaged N200 peak-latency on NDT within the hierarchical drift-diffusion Model 1 for *each* visual noise condition. In addition, the residual NDT in Model 1 were similar across each visual noise types while N200 peak-latency described the differences in NDT across the different visual noise types, a result to be expected if residual NDT reflects motor execution time (MET).

Our findings in this study closely match findings discussed by Thorpe et al. (1996), Martin et al. (2010), Zhang et al. (2016), and Loughnane et al. (2016) as discussed in the Introduction of this paper. In addition, Stanford et al. (2010) showed that when evidence accumulation was blocked for only 100 ms after target onset, monkeys were able to complete their decision, while when evidence accumulation was blocked for 200 ms after target onset, the monkeys were at chance accuracy. The work by Stanford et al. (2010) suggests that visual encoding time must have a latency between 100 ms and 200 ms for a simple color discrimination task. A phenomena of N200 peak-latencies reflecting VET would also explain why N200 latencies have been found to be affected by bottom-up and top-down attention (Vogel and Luck, 2000). In this study we found significant noise-condition effects in N200 peak-latency (see **Table 1**) such that N200 peak-latencies become longer with larger amounts of visual noise contrast (this finding was not observed in the N200 deflection times), and it is expected the VET should become longer with larger amounts of visual noise due to a higher cognitive difficulty of target-noise (e.g. figure-ground) segregation. Furthermore we found some individual differences in N200 peak-latencies across participants, which matches expectations that individual differences in cognitive processing exists that explains individual differences in behavior (e.g. see Schubert et al., 2017, 2018).

### Evidence AGAINST N200 peak-latencies tracking visual encoding time

Single-trial N200 latencies may not be related to VET as indicated by the positive slopes not equal to 1 found between single-trial N200 peak-latencies and NDT in Model 2. However, it is also possible that the single-trial N200 latencies *do* give a satisfactory account of VET but fitting simple drift-diffusion parameter relationships fails on single-trials (see Hawkins et al., 2017), especially with the presence of contaminant processes. This would explain why we found a slope near 1 between single-trial N200 peak-latencies and response times (see bottom right panel of **Figure 6**) but no evidence of this slope of 1 (albeit a significant and positive relationship) when fitting DDM parameters to N200 peak-latencies on single trials. Better theoretical models of behavior with trial-to-trial variability in NDT are probably needed (see Boehm et al., 2018), such as cognitive models that include both decision processes and mind-wandering processes (e.g. see van Vugt et al., 2015; Mittner et al., 2016). Another alternative is a joint model of single-trial N200 latencies and human behavior that can be fit with Bayesian MCMC sampler programs, discussed below.

### A neuro-cognitive theory of decision making

A neuro-cognitive model of encoding, evidence accumulation, and motor response is proposed that is identified with separable measures of VET and MET that are estimated from observed EEG, accuracy, and response time distributions. Fitting this type of model to data would allow direct hypothesis testing of observed trial-to-trial differences and individual differences in neural signals and human behavior during quick decision making.

Neuro-cognitive hierarchical models of VET, evidence accumulation, and motor response time become identifiable when treating EEG measures (e.g. N200 peak-latencies) as measures of VET. The theory is an extension of a specific class of decision-making models (e.g. “drift-diffusion” models, see Ratcliff and McKoon, 2008, for a review) which predicts that response time and accuracy are explained by a continuous accumulation of evidence towards certain pre-decided evidence thresholds (see **Figure 4**).

A drift-diffusion model of accuracy (*w* = 0 or *w* = 1) and response time *t* on one trial is predicted to be a function of VET *τ*_*e*_, MET *τ*_*m*_, the drift rate δ which is the average rate of evidence accumulation within one trial, the diffusion coefficient *ς* which is the variance around the average rate of evidence accumulation within a trial, and the amount of evidence *α_t_* required to make a decision, which may be assumed to decrease with time. The bias towards correct or incorrect responses is assumed to be equal to *z*_0_ = 0.5α_t_. The joint density *f* of RT *t* and accuracy *w* of this simplified diffusion model is given in **Equation 16**. The density is derived from the solution given by Ratcliff (1978) and Tuerlinckx (2004).

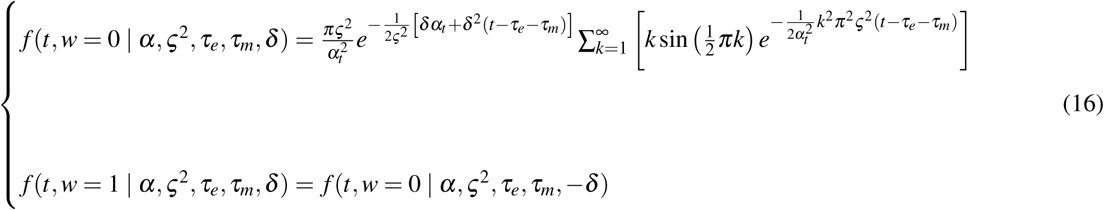

Note that the density in **Equation 16** is classically unidentifiable when estimating the model with only response time and accuracy observations for two reasons. 1) Visual encoding time *τ*_*e*_ and motor execution time *τ*_*m*_ both contribute to non-decision time τ = *τ*_*e*_ + *τ*_*m*_ which approximately equals the smallest observed response times that are not due to contaminant trials. 2) Only two of the three parameters related to evidence accumulation (i.e. drift rate δ, the diffusion coefficient ς, and the boundary separation α) can be found with behavioral data alone. For example, if α_t_ is constant with *t*, multiplying ς by two and dividing both α and δ by two would result in the same fit of choice-RT (Wabersich and Vandekerckhove, 2014). Ideally, a version of the model given in **Equation 16** with four free parameters (e.g. *τ*_*e*_, *τ*_*m*_, d and a constant α in time *t*) would be estimated in JAGS (JASP Team, 2017) or Stan (Carpenter et al., 2016) to allow easy fitting of hierarchical models using the likelihood approximation by Navarro and Fuss (2009) (or Wiecki et al., 2013, for the likelihood approximation extended to intrinsic drift-rate variability). Alternatively, the model may be approximated by shortcuts in JAGS code with hierarchical models; research currently being conducted by the authors of this manuscript.

Fitting neuroscientific results to cognitive models that explain human behavior is a growing field of research (Mulder et al., 2014; Turner et al., 2015; de Hollander et al., 2016; Turner et al., 2017). The goal of future work is to further develop theory that reconciles observed brain function in the Cognitive Neuroscience literature with theoretical behavioral-cognitive relationships. This work could lead to the ability to fit the full density given in **Equation 16** with *identifiable* parameters that explain both psychological phenomena and brain electrophysiology. This would allow researchers and to directly test hypotheses in experimental data and provide researchers and clinicians with a method to explore the cognitive and brain-computational underpinnings behind observed human behavior.

## Acknowledgements

MDN, JV, and RS were supported by NSF grant #1658303. AG was supported by the Summer Undergraduate Research Program at the University of California, Irvine. Siyi Deng and Sam Thorpe are thanked for their contributions to the FEM solutions and anatomical fMRI image generation. Gabriel Weindel is thanked for his early comments and reanalysis of the public data paired with this paper. We would also like to thank Paul L. Nunez for his comments on the spline Laplacian-localization technique used in this paper. Two anonymous reviewers are appreciated for their comments that lead to the improvement of this work. This study extends initial work presented in Chapter 4 of the PhD dissertation by Nunez (2017).

Author Contributions
MDN, JV, and RS designed research. AG and MDN performed research. MDN analyzed data. MDN, AG, JV, and RS wrote the paper.

## Notes

**Conflict of interest statement:** The authors declare no competing financial interests.

## References

Acunzo DJ, MacKenzie G, van Rossum MC (2012) Systematic biases in early ERP and ERF components as a result of high-pass filtering. Journal of neuroscience methods 209:212–218.

Bach M, Meigen T (1992) Electrophysiological correlates of texture segregation in the human visual evoked potential.Vision research 32:417–424.

Boehm U, Annis J, Frank M, Hawkins G, Heathcote A P, Kellen D, Krypotos AM, Lerche V, Logan GD, Palmeri T, Ravenzwaaij Dv, Servant M, Singmann H, Starns J, Voss A, Wiecki T, Matzke D, Wagenmakers EJ (2018) Estimating between-trial variability parameters of the diffusion decision model.

Carpenter B, Gelman A, Hoffman M, Lee D, Goodrich B, Betancourt M, Brubaker MA, Guo J, Li P, Riddell A (2016) Stan: A probabilistic programming language. Journal of Statistical Software 20:1–37.

de Hollander G, Forstmann BU, Brown SD (2016) Different ways of linking behavioral and neural data via computational cognitive models. Biological Psychiatry: Cognitive Neuroscience and Neuroimaging 1:101–109.

Deng S, Winter W, Thorpe S, Srinivasan R (2012) Improved surface laplacian estimates of cortical potential using realistic models of head geometry. IEEE Transactions on Biomedical Engineering 59:2979–2985.

Dickey JM, Lientz BP (1970) The weighted likelihood ratio, sharp hypotheses about chances, the order of a markov chain. The Annals of Mathematical Statistics 41:214–226.

Dmochowski JP, Norcia AM (2015) Cortical components of reaction-time during perceptual decisions in humans. PloS one 10:e0143339.

Drugowitsch J, Moreno-Bote R, Churchland AK, Shadlen MN, Pouget A (2012) The cost of accumulating evidence in perceptual decision making. Journal of Neuroscience 32:3612–3628.

Fischl B, van der Kouwe A, Destrieux C, Halgren E, Ségonne F, Salat DH, Busa E, Seidman LJ, Goldstein J, Kennedy D et al. (2004) Automatically parcellating the human cerebral cortex. Cerebral Cortex 14:11–22.

Fox KCR, Spreng RN, Ellamil M, Andrews-Hanna JR, Christoff K (2015) The wandering brain: Meta-analysis of functional neuroimaging studies of mind-wandering and related spontaneous thought processes. Neuroimage 111:611–621.

Gelman A, Rubin DB (1992) Inference from iterative simulation using multiple sequences. Statistical Science pp. 457–472.

Gold JI, Shadlen MN (2007) The neural basis of decision making. Annual Review of Neuroscience 30:535–574.

Guan M, Vandekerckhove J (2016) A bayesian approach to mitigation of publication bias. Psychonomic Bulletin & Review 23:74–86.

Hawkins GE, Mittner M, Boekel W, Heathcote A, Forstmann BU (2015) Toward a model-based cognitive neuroscience of mind wandering. Neuroscience 310:290–305.

Hawkins GE, Mittner M, Forstmann BU, Heathcote A (2017) On the efficiency of neurally-informed cognitive models to identify latent cognitive states. Journal of Mathematical Psychology 76:142–155.

JASP Team (2017) JASP (Version 0.8.1.2) [Computer software]. Jeffreys H (1961) Theory of probability Oxford University Press.

Kass RE, Raftery AE (1995) Bayes factors. Journal of the American Statistical Association 90:773–795.

Kayser J, Tenke CE (2003) Optimizing PCA methodology for ERP component identification and measurement: theoretical rationale and empirical evaluation. Clinical Neurophysiology 114:2307–2325.

Kayser J, Tenke CE (2015) Issues and considerations for using the scalp surface Laplacian in EEG/ERP research: A tutorial review. International Journal of Psychophysiology 97:189–209.

Kelly SP, O’Connell RG (2015) The neural processes underlying perceptual decision making in humans: recent progress and future directions. Journal of Physiology-Paris 109:27–37.

Lamme VAF (1995) The neurophysiology of figure-ground segregation in primary visual cortex. Journal of Neuroscience 15:1605–1615.

Lamme VAF, Roelfsema PR (2000) The distinct modes of vision offered by feedforward and recurrent processing. Trends in neurosciences 23:571–579.

Lamme VAF, Zipser K, Spekreijse H (2002) Masking interrupts figure-ground signals in v1. Journal of cognitive neuroscience 14:1044–1053.

Lee MD (2008) Three case studies in the Bayesian analysis of cognitive models. Psychonomic Bulletin & Review 15:1–15.

Lee MD, Newell BJ (2011) Using hierarchical Bayesian methods to examine the tools of decision-making. Judgment & Decision Making 6.

Lin CT, Chuang CH, Kerick S, Mullen T, Jung TP, Ko LW, Chen SA, King JT, McDowell K (2016) Mind-wandering tends to occur under low perceptual demands during driving. Scientific Reports 6.

Loughnane GM, Newman DP, Bellgrove MA, Lalor EC, Kelly SP, OConnell RG (2016) Target selection signals influence perceptual decisions by modulating the onset and rate of evidence accumulation. Current Biology 26:496–502.

Luck SJ (2012) Event-related potentials In Cooper H, Camic PM, Long DL, Panter AT, Rindskopf D, Sher KJ, editors, APA Handbook of Research Methods in Psychology: Volume 1, Foundations, Planning, Measures, and Psychometrics. American Psychological Association.

Lui KK, Nunez MD, Cassidy JM, Vandekerckhove J, Cramer SC, Srinivasan R (2018) Timing of readiness potentials reflect a decision-making process in the human brain. bioRxiv 338806.

Makeig S, Bell AJ, Jung TP, Sejnowski TJ (1996) Independent component analysis of electroencephalographic data. Advances in Neural Information Processing Systems 8:145–151.

Martin T, Huxlin KR, Kavcic V (2010) Motion-onset visual evoked potentials predict performance during a global direction discrimination task. Neuropsychologia 48:3563–3572.

Miąsko T (2017) pyjags (version 1.2.2) [computer software] https://github.com/tmiasko/pyjags.

Mishkin M, Ungerleider LG, Macko KA (1983) Object vision and spatial vision: two cortical pathways. Trends in neurosciences 6:414–417.

Mittner M, Hawkins GE, Boekel W, Forstmann BU (2016) A neural model of mind wandering. Trends in cognitive sciences 20:570–578.

Mulder MJ, van Maanen L, Forstmann BU (2014) Perceptual decision neurosciences a model-based review. Neuroscience 277:872–884.

Navarro DJ, Fuss IG (2009) Fast and accurate calculations for first-passage times in wiener diffusion models. Journal of Mathematical Psychology 53:222–230.

Neyman J, Pearson ES (1933) On the problem of the most efficient tests of statistical hypotheses. Philosophical Transactions of the Royal Society of London. Series A, Containing Papers of a Mathematical or Physical Character 231:289–337.

Nunez MD (2017) Refining understanding of human decision making by testing integrated neurocognitive models of EEG, choice and reaction time Ph.D. diss., University of California, Irvine.

Nunez MD, Horton C, Deng S, Winter W, Srinivasan R (2017) artscreenEEG (version 0.3.0) [computer software] https://github.com/mdnunez/artscreenEEG.

Nunez MD, Nunez PL, Srinivasan R (2016) Electroencephalography (EEG): neurophysics, experimental methods, and signal processing In Ombao H, Linquist M, Thompson W, Aston J, editors, Handbook of Neuroimaging Data Analysis, pp. 175–197. Chapman & Hall/CRC.

Nunez MD, Srinivasan R, Vandekerckhove J (2015) Individual differences in attention influence perceptual decision making. Frontiers in Psychology 6:18.

Nunez MD, Vandekerckhove J, Srinivasan R (2017) How attention influences perceptual decision making: Single-trial EEG correlates of drift-diffusion model parameters. Journal of Mathematical Psychology 76, Part B:117–130.

Nunez PL, Nunez MD, Srinivasan R (2018) Multi-scale neural sources of eeg: Genuine, equivalent, and representative. bioRxiv 391318.

Nunez PL, Pilgreen KL (1991) The spline-Laplacian in clinical neurophysiology: a method to improve EEG spatial resolution. Journal of Clinical Neurophysiology 8:397–413.

Nunez PL, Silberstein RB, Cadusch PJ, Wijesinghe RS, Westdorp AF, Srinivasan R (1994) A theoretical and experimental study of high resolution EEG based on surface Laplacians and cortical imaging. Electroencephalography and Clinical Neurophysiology 90:40–57.

Nunez PL, Srinivasan R (2006) Electric Fields of the Brain: The Neurophysics of EEG Oxford University Press.

O’Connell RG, Dockree PM, Kelly SP (2012) A supramodal accumulation-to-bound signal that determines perceptual decisions in humans. Nature Neuroscience 15:1729–1735.

Parra LC, Spence CD, Gerson AD, Sajda P (2005) Recipes for the linear analysis of EEG. Neuroimage 28:326–341.

Philiastides MG, Heekeren HR, Sajda P (2014) Human scalp potentials reflect a mixture of decision-related signals during perceptual choices. The Journal of Neuroscience 34:16877–16889.

Plummer M (2003) JAGS: A program for analysis of Bayesian graphical models using Gibbs sampling In Proceedings of the 3rd International Workshop on Distributed Statistical Computing (DSC 2003), Vienna, Austria.

Pommier J, Renard Y (2005) Getfem++, an open source generic c++ library for finite element methods.

Ratcliff R (1978) A theory of memory retrieval. Psychological Review 85:59.

Ratcliff R, McKoon G (2008) The diffusion decision model: theory and data for two-choice decision tasks. Neural Computation 20:873–922.

Roitman JD, Shadlen MN (2002) Response of neurons in the lateral intraparietal area during a combined visual discrimination reaction time task. Journal of neuroscience 22:9475–9489.

Rosenthal R (1979) The file drawer problem and tolerance for null results. Psychological bulletin 86:638.

Rouder JN, Morey RD (2012) Default bayes factors for model selection in regression. Multivariate Behavioral Research 47:877–903.

Schmolesky MT, Wang Y, Hanes DP, Thompson KG, Leutgeb S, Schall JD, Leventhal AG (1998) Signal timing across the macaque visual system. Journal of neurophysiology 79:3272–3278.

Schubert AL, Hagemann D, Frischkorn GT (2017) Is general intelligence little more than the speed of higher-order processing? Journal of Experimental Psychology: General 146:1498.

Schubert AL, Nunez MD, Hagemann D, Vandekerckhove J (2018) Individual differences in cortical processing speed predict cognitive abilities: a model-based cognitive neuroscience account. bioRxiv 374827.

Servant M, White C, Montagnini A, Burle B (2015) Using Covert Response Activation to Test Latent Assumptions of Formal Decision-Making Models in Humans. Journal of Neuroscience 35:10371–10385.

Servant M, White C, Montagnini A, Burle B (2016) Linking theoretical decision-making mechanisms in the simon task with electrophysiological data: A model-based neuroscience study in humans. Journal of Cognitive Neuroscience 28:1501–1521.

Shadlen MN, Kiani R (2013) Decision making as a window on cognition. Neuron 80:791–806.

Stanford TR, Shankar S, Massoglia DP, Costello MG, Salinas E (2010) Perceptual decision making in less than 30 milliseconds. Nature neuroscience 13:379–385.

Sterling TD (1959) Publication decisions and their possible effects on inferences drawn from tests of significanceor vice versa. Journal of the American statistical association 54:30–34.

Straube S, Grimsen C, Fahle M (2010) Electrophysiological correlates of figure–ground segregation directly reflect perceptual saliency. Vision Research 50:509–521.

Thorpe S, Fize D, Marlot C (1996) Speed of processing in the human visual system. Nature 381:520.

Tuerlinckx F (2004) The efficient computation of the cumulative distribution and probability density functions in the diffusion model. Behavior Research Methods, Instruments, & Computers 36:702–716.

Turner BM, Forstmann BU, Love BC, Palmeri TJ, Van Maanen L (2017) Approaches to analysis in model-based cognitive neuroscience. Journal of Mathematical Psychology 76:65–79.

Turner BM, van Maanen L, Forstmann BU (2015) Informing cognitive abstractions through neuroimaging: The neural drift diffusion model. Psychological Review 122:312–336.

van Vugt MK, Taatgen NA, Sackur J, Bastian M (2015) Modeling mind-wandering: a tool to better understand distraction In Proceedings of the 13th International Conference on Cognitive Modeling, pp. 252–257. University of Groningen.

Vandekerckhove J, Tuerlinckx F (2008) Diffusion model analysis with MATLAB: A DMAT primer. Behavior Research Methods 40:61–72.

Vandekerckhove J, Tuerlinckx F, Lee MD (2011) Hierarchical diffusion models for two-choice response times. Psychological Methods 16:44.

Vanrullen R, Thorpe SJ (2001) The time course of visual processing: from early perception to decision-making. Journal of cognitive neuroscience 13:454–461.

Verdinelli I, Wasserman L (1995) Computing Bayes factors using a generalization of the savage-dickey density ratio. Journal of the American Statistical Association 90:614–618 00422.

Vogel EK, Luck SJ (2000) The visual N1 component as an index of a discrimination process. Psychophysiology 37:190–203.

Voss A, Rothermund K, Voss J (2004) Interpreting the parameters of the diffusion model: An empirical validation. Memory & Cognition 32:1206–1220.

Wabersich D, Vandekerckhove J (2014) Extending JAGS: A tutorial on adding custom distributions to JAGS (with a diffusion model example). Behavior Research Methods 46:15–28.

Wagenmakers EJ (2009) Methodological and empirical developments for the Ratcliff diffusion model of response times and accuracy. European Journal of Cognitive Psychology 21:641–671.

Wagenmakers EJ, Lodewyckx T, Kuriyal H, Grasman R (2010) Bayesian hypothesis testing for psychologists: A tutorial on the savage–dickey method. Cognitive psychology 60:158–189.

Webster MA, De Valois RL et al. (1985) Relationship between spatial-frequency and orientation tuning of striate-cortex cells. JOSA A 2:1124–1132.

Wiecki TV, Sofer I, Frank MJ (2013) HDDM: hierarchical bayesian estimation of the drift-diffusion model in python. Frontiers in neuroinformatics 7:14.

Zhang Q, van Vugt M, Borst JP, Anderson JR (2018) Mapping working memory retrieval in space and in time: A combined electroencephalography and electrocorticography approach. NeuroImage.

Zhang Q, Walsh MM, Anderson JR (2016) The effects of probe similarity on retrieval and comparison processes in associative recognition. Journal of Cognitive Neuroscience.

